# Comparative developmental genomics of sex-biased gene expression in early embryogenesis across mammals

**DOI:** 10.1101/2022.03.17.484606

**Authors:** Victorya Richardson, Kai Zhang, Nora Engel, Rob J Kulathinal

## Abstract

Mammalian gonadal sex is determined by the presence or absence of a Y chromosome. In males, the Y chromosome initiates male gonadogenesis and the subsequent production of male-specific hormones defines the male state of each cell in the organism. In females, the lack of a Y chromosome and the presence of two X chromosomes triggers the development of female gonads, hormones, and cellular identity. However, sex chromosome-linked genes encoding dosage-sensitive transcription and epigenetic factors are expressed well before gonad formation and have the potential to establish sex-biased expression. Here, we apply a comparative bioinformatics analysis on published single-cell datasets from mouse and human during very early embryogenesis–from two-cell to preimplantation stages–to characterize sex-specific signals and to assess the degree of conservation among early-acting sex-specific genes and pathways. Clustering and regression analyses of gene expression across samples reveal that sex initially plays a significant role in overall gene expression patterns at the earliest stages of embryogenesis. In addition, gene expression signals from male and female gametes during fertilization may still be present. Although these transcriptional sex effects rapidly diminish, the sex-biased expression of epigenetic enzymes has the potential to establish sex-specific patterns that persist beyond preimplantation. Sex-biased genes appear to form sex-specific protein-protein interaction networks across preimplantation stages in both mammals. While the distribution of sex-differentially expressed genes (sexDEGs) in early embryonic stages are similar in mice and humans, the genes involved are generally different. Non-negative matrix factorization (NMF) on male and female transcriptomes generated clusters of genes with similar expression patterns across sex and developmental stages including post-fertilization, epigenetic, and preimplantation ontologies conserved between mouse and human. This comparative study uncovers much earlier than expected sex-specific signals in mouse and human embryos that pre-date hormonal signaling from the gonads. These early signals are diverged with respect to orthologs yet conserved in terms of function with important implications in the use of genetic models for sex-specific disease.

## INTRODUCTION

The dichotomy between the sexes is distinct, pervasive, and often extreme across the eukaryotic tree of life^1-3^. Males and females harbor both conspicuous and cryptic differences in reproduction, physiology, morphology, and behavior despite sharing the same allelic variation. Sexual dimorphism is thought to have originated over a billion years ago, representing a defining hallmark of eukaryote diversity, particularly among animals. Indeed, the ubiquity of sexual dimorphism represents a conserved biological trait shared across the diversity of life. Evolutionary mechanisms such as sexual selection^1^ and sexual conflict^4^ have been hypothesized to maintain and promote the presence of sexually dimorphic traits, most of which are taxon-specific.

While sex-specific differences are readily observed in adults, these differences are initiated by early acting sex determining and compensatory mechanisms that occur early in embryogenesis. In mammals, the process of establishing sex differences has traditionally been divided into two phases: i) an initial genetic stage of gonad formation, also known as “sex determination”, with SRY as the master regulator of male gonadogenesis and ii) a later secondary stage of sexual differentiation regulated by sex hormones^5^.

However, this view ignores the consequences of the inherent differences in sex chromosome composition (i.e., XX in females and XY in males) which affect the embryo beginning soon after fertilization, and effects on gonadal and non-gonadal tissues throughout the organism’s lifespan^6^. Sex-linked genes, some of which are transcription and epigenetic factors with downstream autosomal targets, are expressed early in embryogenesis^7-11^. In addition, female embryos undergo X chromosome inactivation during implantation, a massive epigenetic event that has been hypothesized to alter the levels of epigenetic factors in female cells relative to male cells^12-17^. These early differences in regulatory factors have the potential to modify the transcriptional and epigenetic landscape in a sex-specific manner that can persist across the lifespan.

Recently, developmental studies reported transcriptional programs using single-cell RNA-sequencing experiments in early embryogenesis^18-20^. Yet sex-biased gene expression has rarely been surveyed during or preceding preimplantation which encompasses the stages from zygote to blastocyst before the embryo interacts and connects with the uterus^21,22^. The few studies that exist show that sex-biased gene expression from both sex chromosomes and autosomes is detectable in pre-implantation embryos and in embryonic stem cells in rodents, bovine and primates, including humans^7,8,11^. These differences can propagate through regulatory networks, resulting in distinct male and female cell states well before gonadal development.

In this paper, we identify and compare male and female gene expression levels in preimplantation embryogenesis across two mammalian species using published transcriptomic time series. Our goals are twofold: i) to identify early sex-specific patterns of gene expression in the earliest stages of embryogenesis in both mouse and human and ii) to compare how conserved early acting sex-specific networks are across mammalian lineages. We identify very early acting sex-specific genes and networks in both mice and humans that, surprisingly, are not shared among orthologs. In contrast, however, functional ontologies appear to be similar between mice and humans across developmental stages. Our study provides support for a dynamic stage- and sex-specific landscape of gene expression that underlies conserved phenotypes of development and sexual identity in the earliest stages of an individual’s lifecycle.

## MATERIALS AND METHODS

### Sequence retrieval, pre-processing, and normalization

Single-cell transcriptomic data were downloaded from two independent studies surveying the earliest stages of embryonic development in mouse and human: 106 mouse samples representing 2-cell to early blastocyst development^9^ (https://www.ncbi.nlm.nih.gov/geo/query/acc.cgi?acc=GSE80810) and 1,529 human samples stemming from 8-cell to late blastocyst^18^ (https://www.ebi.ac.uk/arrayexpress/experiments/E-MTAB-3929/). A summary of the temporal sample sources for the pair of datasets is found in Supplementary Table S1. Mouse genes with RIKEN annotations (n=1,860) were removed from the dataset. Lowly expressed genes and samples with poor coverage were filtered using R package “Seurat” (min.cells=3, min.features=350)^23-26^. Inter-cell normalization was performed using a deconvolution approach for single-cell transcriptomic data with many zero values^27^ (Supplementary Figure S1).

### Sexing single cell data

Single cells were sexed by determining the expression ratios of two early expressed genes, *Xist* and *Eif2s3y*, located on respectively the X and Y chromosomes. *Xist* to *Eif2s3y* ratios were estimated individually on a cell-by-cell basis with samples harboring ratios of at least 1.5 considered as female and ratios below 1 were taken as male. Samples with ambiguous ratios between 1 and 1.5 were removed from the analysis.

### Sex-differential expression analyses

Sex-differentially expressed genes between male and female cells (sexDEGs) were independently identified at each embryonic stage using “DESeq2” in R, with default parameters. To contrast expression levels between males and females at each stage, we used the design formula “∼sex_stage”, where “sex_stage” is a column of combined sex and stage data. Log_2_ fold change results were transformed using lfcShrink() in R with default parameters. Genes with |log_2_ fold change| ≥ 0.58 and adjusted p-values < 0.05 were marked as differentially expressed. Results of the analyses are reported in Supplementary Tables S2 and S3 for mouse and human, respectively.

### Non-negative matrix factorization

To identify longitudinal sex-specific subnetworks of co-expression, count data from the mouse and human datasets were divided into male- and female-specific count tables (mouse samples: 71 female, 35 male; human samples: 821 female, 708 male). The resulting subsets were filtered for low expression (min.cells = 3, min.features = 350) and log-normalized using R package “Seurat”^24^. To identify the optimal matrix rank (or number of resulting clusters), NMF was run 10 times for each user-inputted rank across ranks 5 to 30, using R package “NMF”^28^. For each sex-specific count table, the rank used for identification of clusters (or metagenes) was chosen as the rank corresponding to the highest cophenetic correlation coefficient (see supplementary cophenetic correlation plot). The coefficient, or *W*, matrix for the chosen rank was extracted for subsequent gene set enrichment analysis (Supplementary Tables S8-S11).

### Functional enrichment analysis

Gene collections (ontology gene sets, hallmark gene sets, and regulatory target gene sets) were accessed from the Molecular Signatures Database (MSigDB)^29,30^. Pre-ranked gene set enrichment analysis was performed on each metagene using R package “fgsea” (minSize = 15, maxSize = 500, scoreType = ‘pos’), with genes ranked according to their entry in the coefficient matrix corresponding to the metagene column^31^. Significant gene pathways (adjusted p-value < 0.05) for each metagene were kept for comparison of enrichment of sex-specific emergent subnetworks between mice and humans (Supplementary Tables S4-S7).

## RESULTS

### Transcriptomics of early development in mouse and human

Nearly half of all autosomal genes are expressed in early embryogenesis, i.e., during their first week of development (Figure 1A), in both sexes of mouse and human (Figure 1B). While the number of expressed autosomal genes remains relatively constant from the two-cell to pre-implantation stages across sex and species, a larger variance in the fraction of sex-chromosomal genes expressed across mouse and human developmental stages is observed. The ratio of X:A gene expression is different between mammals with relatively lower fractions of X-linked genes being expressed in mouse. In addition, a larger proportion of Y-chromosome genes are expressed in male human compared to male mouse. However, the Y-linked expressed fraction is generally much lower than the expressed fraction of X-linked and autosomal genes in both mammals (Figure 1B).

**Figure 1.**
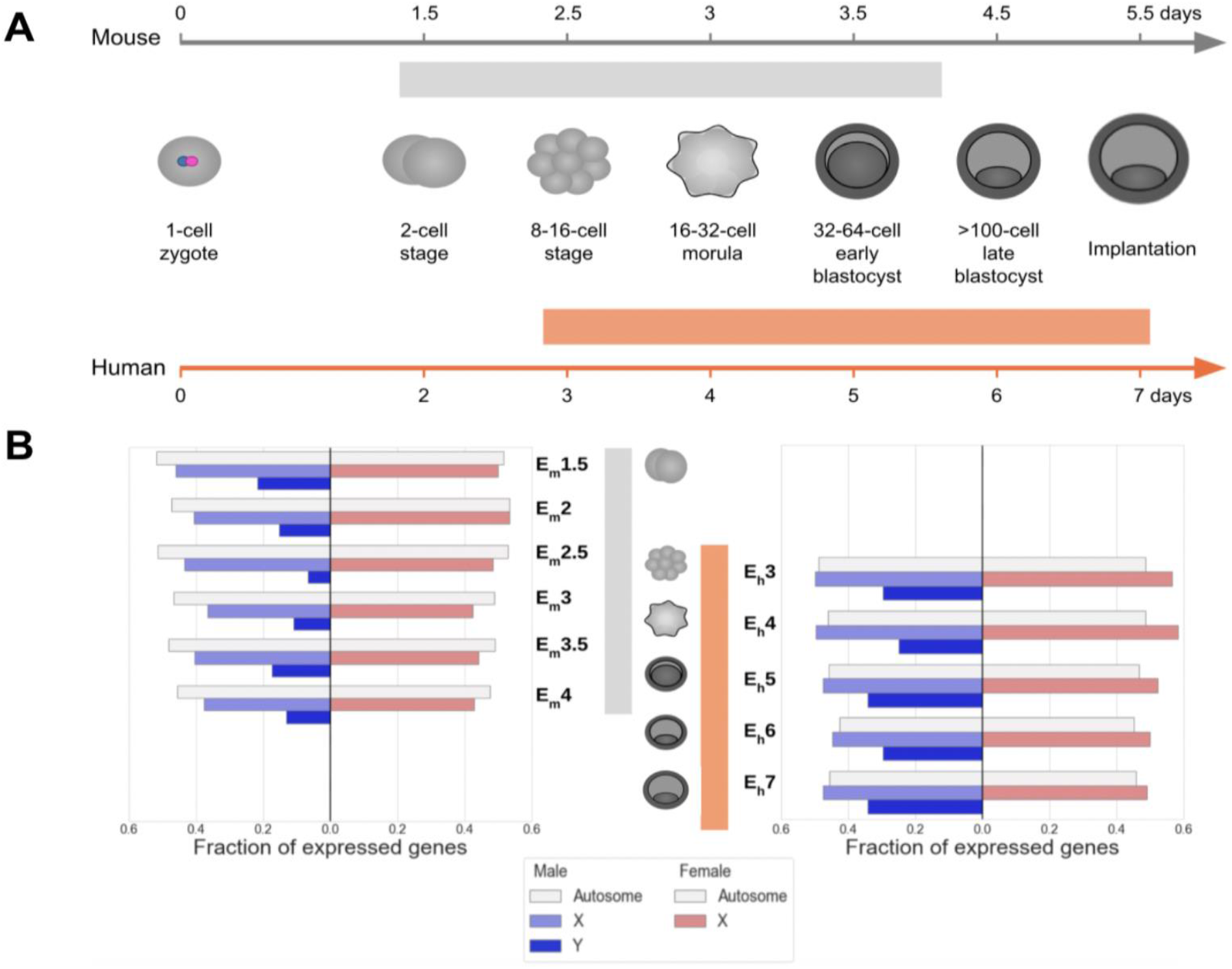
Developmental timeline and gene expression levels between mouse and human samples used in this study. (A) The relative timing of embryogenic stages from zygote to post-implantation differs across the two mammals, mouse (gray) and human (orange). Samples taken from a pair of published datasets derived from mouse and human represent a series of very early developmental stages (Borensztein et al. 2016; Petropoulos et al. 2016). (B) Fraction of genes expressed that are found on the autosomes, X-, and Y-chromosomes in males (blue) and females (red) are displayed separately for mouse (left) and human (right) datasets.

Of the 1,000 most highly expressed genes in mouse preimplantation stages, the number of genes that encode transcription factors (TF) and epigenetic enzymes (EE) peaked at the two-cell stage with steadily diminishing numbers as development proceeded (E_m_1.5: 51 TFs, 27 EEs; E_m_3: 24 TFs, 12 EEs). A similar trend was observed for human embryos in transcription factors (E_m_3: 49 TFs; E_m_7: 23 TFs). However, the number of expressed genes encoding EEs was almost constant between stages. Surprisingly, the orthologs of only 17 TFs and 6 EEs expressed in the mouse were detected in human embryos. Among the common regulatory factors between mouse and human, most were expressed at analogous stages of early embryonic development, such as *Atf4, Elf3, Sall4, Tfap2c, Hdac1, Kdm5b* and *Tet1*.

Several the so-called “pluripotency factors”^32,33^, which include transcription factors, epigenetic enzymes, and signaling molecules, were detected in the two datasets with several uniquely expressed in one of the two species. For example, *Dnmt3b, Dnmt3l, Sall4*, and *Tead4* were present in both mouse and human while *Nanog, Esrrb, Gata4*, and *Pou5f1* (*Oct4*) were not detected in human embryos, and *Klf4, Myc and Dnmt3a* were not detected in mouse.

### Relative contribution of developmental stage vs. sex to gene expression in early embryogenesis

Normalized genome-wide expression counts from each sample were found to primarily cluster according to developmental stage progression in both mouse and human (Figure 2A-B). The first two principal components explained 63% and 50% of the total variance in gene expression in, respectively, mouse and human. While male and female samples from each developmental stage clustered in a time-dependent manner, it was difficult to visually differentiate among sexed samples via PCA. Male and female samples appear to weakly cluster together at very early stages and less during later stages of pre-implantation embryogenesis. To better understand the quantitative contribution of sex across development stages, we employed a linear regression model and found that sex explained nearly a quarter of the genetic variance in gene expression during the earliest stages of embryogenesis in both mouse and human (Figure 2C-D) but that this contribution of sex rapidly decreased. This rapid diminution in the expression variation that is explained by sex reflects that sex’s relative role decreases rapidly and substantially across very early development. Genes with the highest and lowest principal component scores for the top 30 principal components of gene expression data in mice and humans are shown in Supplementary Figures S2 and S3.

**Figure 2.**
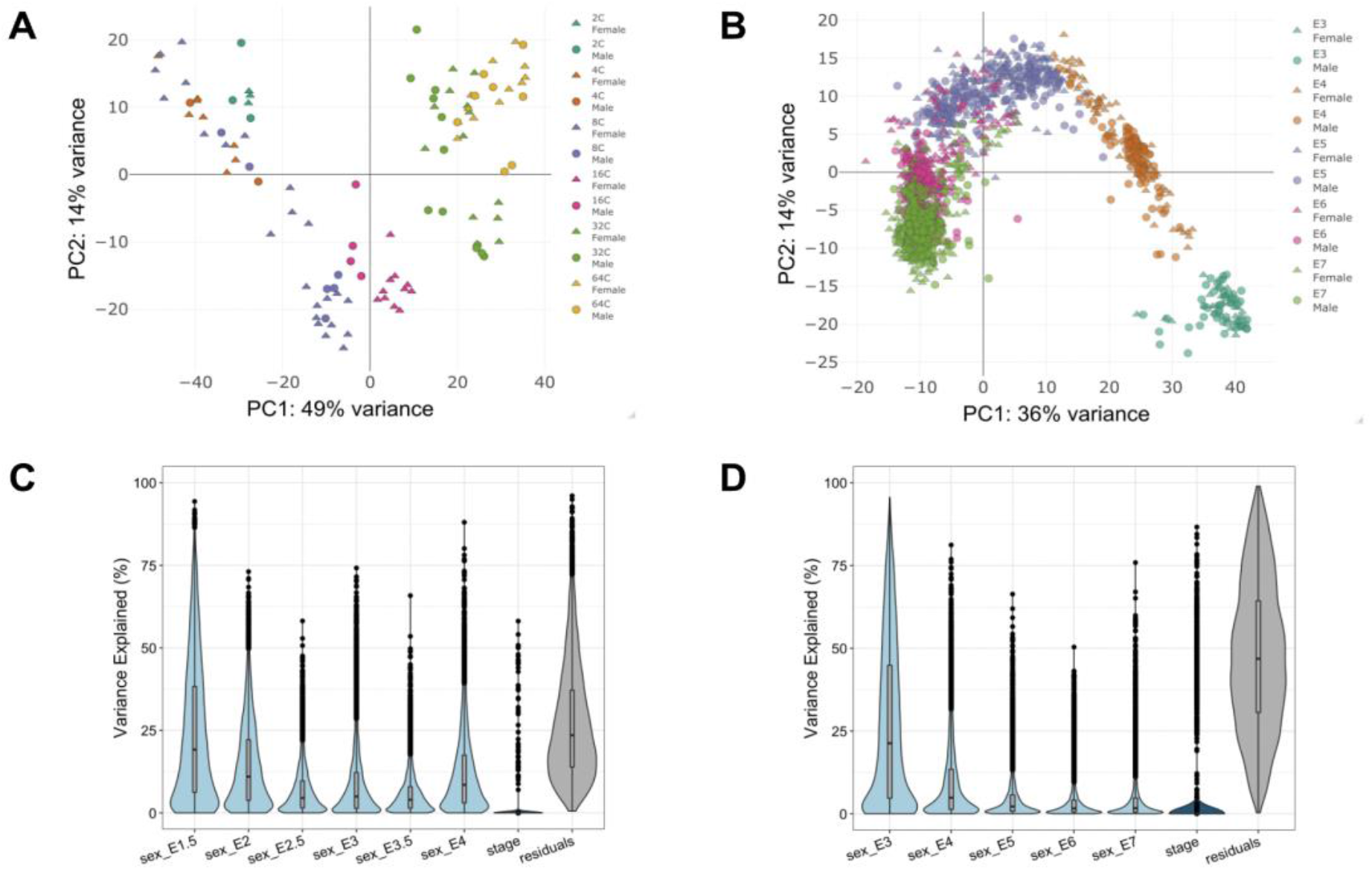
Contribution of embryonic stage and sex in total gene expression variation across very early mouse and human embryogenesis. Principal component analysis of (A) mouse embryonic samples (n=106) and (B) human embryonic samples (n=1,529). Normalized data were transformed using the *varianceStabilizingTransformation* function in DESeq2. In both mice and humans, the majority of the total variance is explained by PC1 and PC2. The distribution and density of the fraction in expression variance for each gene that can be explained by embryonic stage and sex in (C) mouse and (D) human.

### Enrichment of sex-differentially expressed genes

Common enrichment patterns among sex-differentially expressed genes (sexDEGs) are observed in males and females in both mouse and human (Table 1). As expected due to the different number of sex chromosomes in males and females, sexDEGs that are X- and Y-linked are over-enriched in, respectively, females and males across both species (Table 1). In contrast, the number of sex-differentially expressed autosomal genes is statistically under-represented in both sexes in most early developmental stages in mouse and human. X-linked genes also have lower than expected number of sexDEGs in early stages (though with low sample sizes). However, the X-chromosome becomes over-enriched for sexDEGs during the blastocyst stage. In terms of functional annotations, while the majority of DEGs are protein-coding genes, these genes are under-enriched for DEGs in many developmental stages with a greater effect in humans, likely due to increased samples and statistical power (Table 1). Similarly, this lack of power likely impedes our ability to detect enrichment in epigenetic and transcription factor sexDEGs in early embryogenesis.

**Table 1.** Enrichment of differentially expressed genes between males and females (sexDEGs) for function and genomic location. For each stage in (A) mouse and (B) human, we tested for enrichment (over and under) of sex differentially expressed genes (sexDEGs) according to general functional classes (protein-coding, epigenetic factors, transcription factors) and chromosomal location. P-values shaded in red represent an under-enrichment while p-values shaded in green represent an over-enrichment. P-values with asterisk are low-confidence due to low sample size in the given group.

Male
|  | Protein coding |  | Epigenetic Factors |  | Transcription Factors |  | X Chromosome |  | Y chromosome |  | Autosome |  |
| --- | --- | --- | --- | --- | --- | --- | --- | --- | --- | --- | --- | --- |
|  | Odds ratio | P Value | Odds ratio | P Value | Odds ratio | P Value | Odds ratio | P Value | Odds ratio | P Value | Odds ratio | P Value |
| **E1.5** | 1.523 | 0.525* | 0.841 | 1.000* | 0.887 | 1.000* | 0.000 | 0.177* | 45.136 | 0.002* | 2.139 | 0.289 |
| **E2** | 0.728 | 0.272 | 0.869 | 1.000* | 1.274 | 0.401 | 0.469 | 0.448* | 68.316 | << 0.01* | 0.696 | 0.201 |
| **E2.5** | 0.822 | 0.657* | 0.000 | 0.414* | 1.075 | 0.848 | 0.613 | 0.774* | N/A | << 0.01* | 0.880 | 0.711 |
| **E3** | 0.383 | << 0.01 | 0.716 | 0.388* | 0.750 | 0.067 | 0.887 | 0.668 | 40.439 | << 0.01* | 0.676 | 0.002 |
| **E3.5** | 1.229 | 0.738* | 1.240 | 0.737* | 1.076 | 0.746 | 1.565 | 0.229* | 130.829 | << 0.01* | 0.586 | 0.041 |
| **E4** | 0.578 | 0.002 | 0.877 | 1.000* | 0.976 | 1.000 | 0.729 | 0.455* | N/A | << 0.01* | 0.560 | << 0.01 |

Female
|  | Protein coding |  | Epigenetic Factors |  | Transcription Factors |  | X Chromosome |  | Y chromosome |  | Autosome |  |
| --- | --- | --- | --- | --- | --- | --- | --- | --- | --- | --- | --- | --- |
|  | Odds ratio | P Value | Odds ratio | P Value | Odds ratio | P Value | Odds ratio | P Value | Odds ratio | P Value | Odds ratio | P Value |
| **E1.5** | 0.680 | 0.257 | 0.656 | 1.000* | 0.813 | 0.846 | 0.732 | 0.800 | N/A | 1.000* | 0.665 | 0.201 |
| **E2** | 1.122 | 0.725 | 0.782 | 1.000* | 1.055 | 0.805 | 2.339 | << 0.01 | 0.000 | 1.000* | 0.562 | << 0.01 |
| **E2.5** | 0.508 | << 0.01 | 1.175 | 0.588* | 1.060 | 0.788 | 10.452 | << 0.01 | 0.000 | 1.000* | 0.166 | << 0.01 |
| **E3** | 0.811 | 0.168 | 0.993 | 1.000* | 0.904 | 0.632 | 2.213 | << 0.01 | N/A | 1.000* | 0.525 | << 0.01 |
| **E3.5** | 0.417 | << 0.01 | 0.278 | 0.050* | 1.030 | 0.848 | 2.639 | << 0.01 | N/A | 1.000* | 0.410 | << 0.01 |
| **E4** | 1.070 | 0.832 | 0.671 | 0.668* | 1.356 | 0.098 | 2.195 | << 0.01 | 0.000 | 1.000* | 0.653 | << 0.01 |

B. HUMAN Male
|  | Protein coding |  | Epigenetic Factors |  | Transcription Factors |  | X Chromosome |  | Y chromosome |  | Autosome |  |
| --- | --- | --- | --- | --- | --- | --- | --- | --- | --- | --- | --- | --- |
|  | Odds ratio | P Value | Odds ratio | P Value | Odds ratio | P Value | Odds ratio | P Value | Odds ratio | P Value | Odds ratio | P Value |
| **E3** | 1.203 | 0.304 | 1.281 | 0.601* | 0.780 | 0.263 | 0.176 | << 0.01* | 252.571 | << 0.01* | 1.135 | 0.509 |
| **E4** | 0.395 | << 0.01 | 1.603 | 0.253* | 0.871 | 0.597 | 0.110 | << 0.01* | 438.699 | << 0.01* | 0.675 | 0.026 |
| **E5** | 0.784 | 0.034 | 0.949 | 1.000* | 0.909 | 0.515 | 0.195 | << 0.01* | 178.027 | << 0.01* | 1.033 | 0.855 |
| **E6** | 0.475 | << 0.01 | 0.509 | 0.255* | 1.093 | 0.509 | 0.841 | 0.628 | 218.255 | << 0.01* | 0.595 | << 0.01 |
| **E7** | 0.539 | << 0.01 | 0.557 | 0.497* | 0.693 | 0.072 | 1.679 | 0.034 | 321.327 | << 0.01 | 0.475 | << 0.01 |

Female
|  | Protein coding |  | Epigenetic Factors |  | Transcription Factors |  | X Chromosome |  | Y chromosome |  | Autosome |  |
| --- | --- | --- | --- | --- | --- | --- | --- | --- | --- | --- | --- | --- |
|  | Odds ratio | P Value | Odds ratio | P Value | Odds ratio | P Value | Odds ratio | P Value | Odds ratio | P Value | Odds ratio | P Value |
| **E3** | 0.701 | 0.022 | 0.484 | 0.444* | 0.455 | 0.001* | 8.710 | << 0.01 | 0.000 | 1.000* | 0.281 | << 0.01 |
| **E4** | 1.263 | 0.053 | 1.297 | 0.394* | 0.820 | 0.164 | 14.919 | << 0.01 | 0.000 | 1.000* | 0.252 | << 0.01 |
| **E5** | 0.427 | << 0.01 | 1.198 | 0.497* | 0.942 | 0.692 | 20.070 | << 0.01 | N/A | 0.070* | 0.161 | << 0.01 |
| **E6** | 0.570 | << 0.01 | 0.632 | 0.215* | 1.139 | 0.216 | 7.291 | << 0.01 | N/A | 0.111* | 0.365 | << 0.01 |
| **E7** | 0.530 | << 0.01 | 1.085 | 0.721* | 1.033 | 0.782 | 6.970 | << 0.01 | N/A | 0.061* | 0.359 | << 0.01 |

### Characterization of sex-specific differences in early mammalian embryogenesis

During the early stages of embryogenesis in both mouse and human, more genes appear to be expressed than not expressed (Figure 3A). However, this ratio quickly becomes approximately 1:1. The number of sex-differentially expressed genes in both mouse and human is small (Figure 3A). In mouse, the total number of sexDEGs increases starting from the four-cell stage and peaks at the sixteen-cell stage (E_m_3). Although the number male-biased genes is eight times higher than female-biased genes at the four-cell and eight-cell stages, this ratio changes at the sixteen-cell stage in which up-regulated female-biased genes are about twice the number of upregulated male DEGs. Our functional enrichment analyses of DEGs found that transcription factors (TFs) were enriched at the four-cell, eight-cell, and 64-cell stages while genes on the sex (X,Y) chromosomes were enriched at the eight-cell and 64-cell stages. None of the stages revealed an enrichment for miRNAs among DEGs. We also found a significant difference in the number of DEGs encoding transcription factors (TFs) over-expressed in females compared with TFs that were over-expressed in males across developmental stages. The number of female-biased TF DEGs follows the same pattern as total DEGs, while the number of male-biased TF DEGs remains constant across developmental stages. In human, a comparable magnitude of sexDEGs is seen across stages with a peak in the late blastocyst stage, where female DEGs were found to be twice as high as male DEGs (Figure 3A).

**Figure 3.**
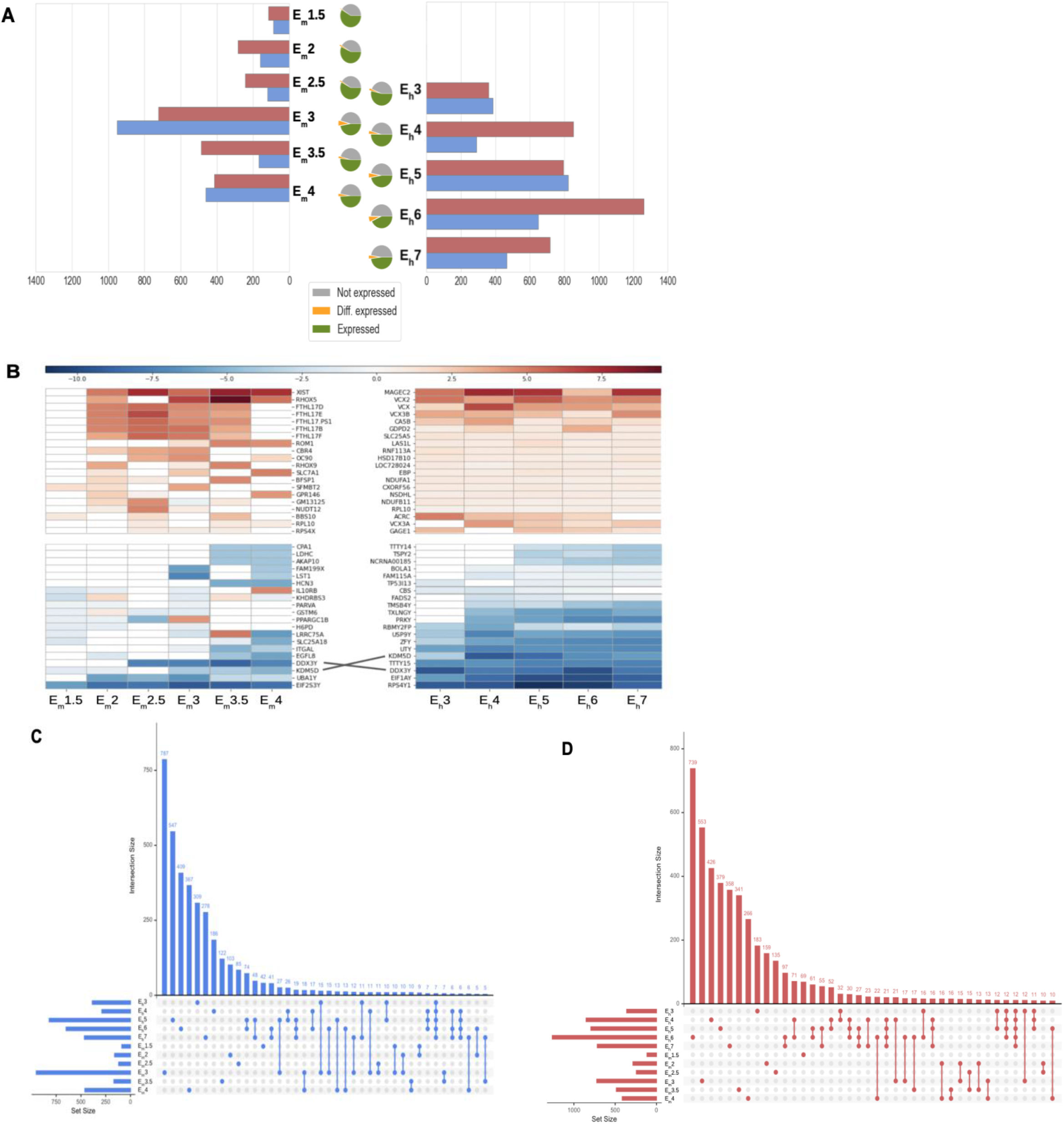
Sex-differentially expressed (sex-biased) genes across early embryonic stages of mouse and human. (A) The number and direction (female-vs male-biased) of sexually differentially expressed genes (sexDEGs) across sampled embryonic stages in mouse and human. The number of sex-biased genes (orange) is observed to be relatively small compared to the total number of expressed genes (green) or non-expressed genes (gray), as seen in the pie-charts. (B) Heatmap of the twenty most female-biased genes (top, red) and the twenty most male-biased genes (bottom, blue) for mouse (left) and human (right). Color intensity indicates log-fold change obtained from a Wald test for differential expression. A pair of orthologs found among these highly sex-biased genes in mouse and human are connected by a line. Upset plots of shared sexDEGs across mouse and human developmental stages for (C) male-biased and (D) female-biased sexDEGs.

Figure 3B presents heatmaps of the top twenty most consistently female-biased (red, top half) and male-biased (blue, bottom half) genes across development stages for mouse (left heatmap) and human (right heatmap). Interestingly, there were only two orthologs between mouse and human that showed similar sex-biased patterns, *KDM5D* and *DDX3Y*, both Y-linked genes. *KDM5D* is a histone lysine demethylase with a repressive transcriptional role and *DDX3Y* is a DEAD-box RNA helicase.

Functional enrichment of female selected clusters shows “nucleobase-containing small molecule metabolic process” involved in processing of nucleotides. However, this analysis for selected clusters of male samples shows metabolic and catabolic processes. (The samples from each stage are shown in different colors.) Surprisingly, the resulting network from male samples was very small with only 51 nodes; however, the female samples resulted in a network with about 4,000 nodes. Almost all the important nodes in each network (based on different centrality criteria) are available in the DEGs list (networks not shown here).

### Differences in sex-differentially expressed protein interactions between mouse and human

In early mouse development, protein-protein interactions are dominated by non-differentially expressed genes (Figure 4A). In the 16-cell stage, we observe a burst of male-specific interactions and in stage 32C we see a similar burst in female-specific interactions while the number of interactions between non-DEGs decreases accordingly. In these “bursts”, interactions between male or female DEGs with non-DEGs also increases, but the same decrease is seen in both male and female in the blastocyst stage. A different pattern is seen in humans. Comparing the normalized fraction of interactions to total DEGs, we don’t see a burst in male- or female-specific interactions like we do in mice but there are still some patterns. Non-DEGs interactions are higher in earlier stages, decrease in middle stages E5 and E6, and increase again in the latest stage. We also observe an increase in female-female DEG interactions in E6, but not like the sharp increase observed in mice. It should be noted that the network sizes for human tend to be smaller here due to fewer human genes mapping to the PPI data.

**Figure 4.**
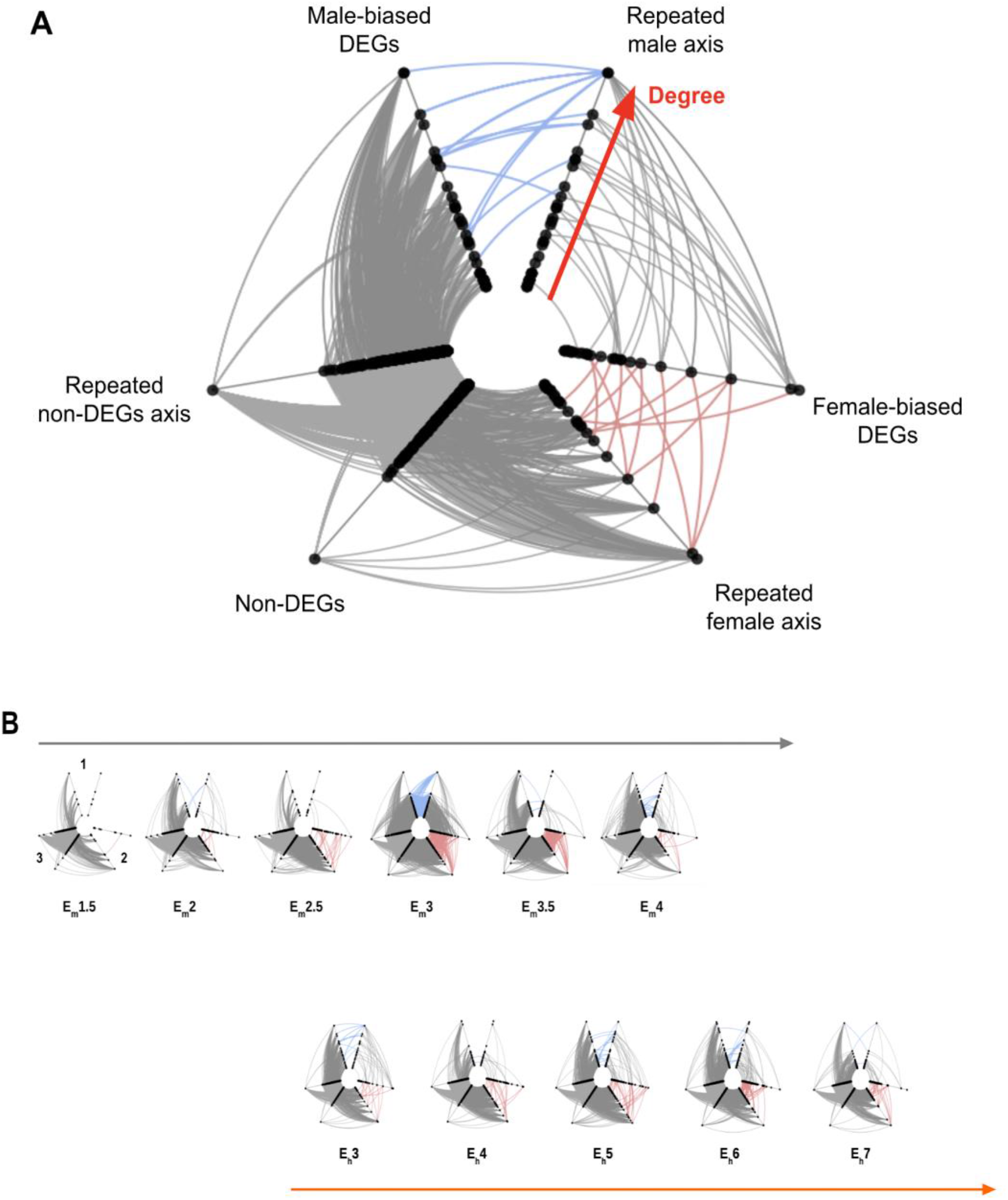
Functional enrichment of emerging sex-biased protein-protein interaction networks across early development stages in mouse and human. (A) Differentially expressed genes between sexes (sexDEGs; male-biased: blue; female-biased: red) are mapped onto full protein-protein interaction networks constructed from expressed genes across embryonic stage. Axes correspond to (i) male-biased DEGs, (ii) female-biased DEGs, and (iii) non-DEGs.

### Lack of conservation signal in orthologous genes across very early mouse and human embryonic stages

A clustering analysis (UMAP) of integrated gene expression data of all samples from both mammals allow us to align both datasets and compare the structure of clusters between mouse and human across timepoints. We observe similar stages in mouse and human in the same region of the graph, though again, it’s difficult to directly compare with the limited mouse samples. Cells are colored based on a general stage of developmental timepoints (see key in Figure 5A) that include enough samples from both mouse and human. Categorizing the cells between mouse and human in this manner is essential for our analysis in Figure 5B-E, which identifies genes that are conserved between mouse and human specifically in each general stage of S3, etc. Therefore, it’s essential to have at least some cells from both species that are in each timepoint to do this comparison.

**Figure 5.**
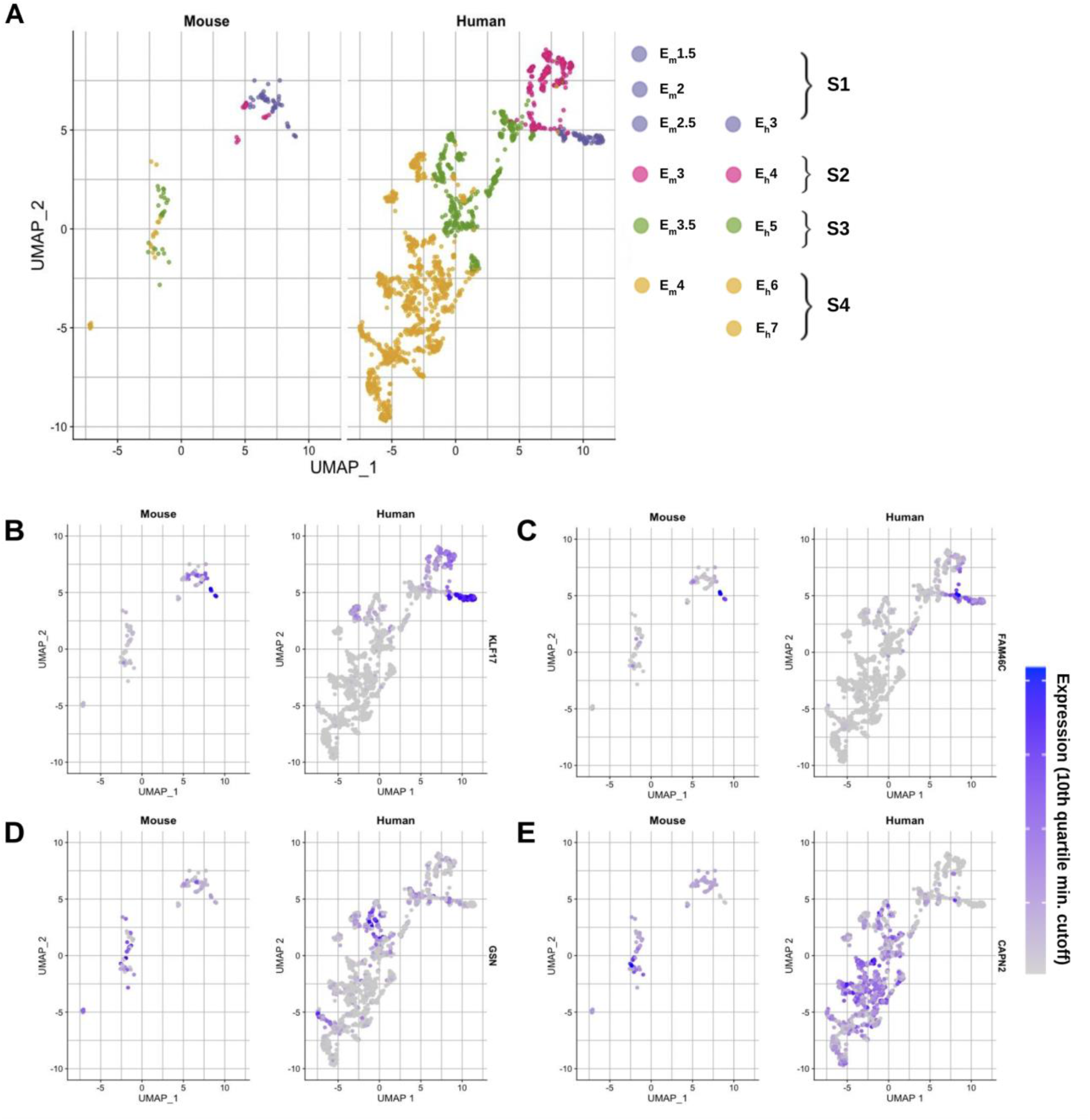
Lack of conservation signal in orthologous genes across very early mouse and human embryonic stages. (A) Integrated UMAP analysis of human and mouse data across embryonic stages. The two datasets were integrated using Seurat to identify common cell types. (B) Expression of the top 8 conserved markers between mouse and human. Expression values were filtered to be within the 10th quantile of the integrated expression data.

In Figure 5B-E, the same UMAPs, but normalized expression levels are highlighted on a gradient in each plot. An example of a highly conserved gene for each stage described in figure 5A (by p-value, see table of top conserved genes for each general timepoint S1, S2, S3, S4). Only genes in the top 10th quartile for expression level in each species are used in the color bar scale; the rest are gray to display contrast of high expression in certain developmental timepoints vs. others.

### Conserved sex-biased genes and networks between mouse and human

We applied non-negative matrix factorization to identify co-expressed genes across stages separately in i) male mouse (Figure 6A), ii) male human (Figure 6B), iii) female mouse (Figure 6C), and iv) female human (Figure 6D). NMF analysis of gene expression data gives unsupervised clusters of genes with similar expression patterns across timepoints (Figure 6A-D) (Supplementary Figures S4-S12).

**Figure 6.**
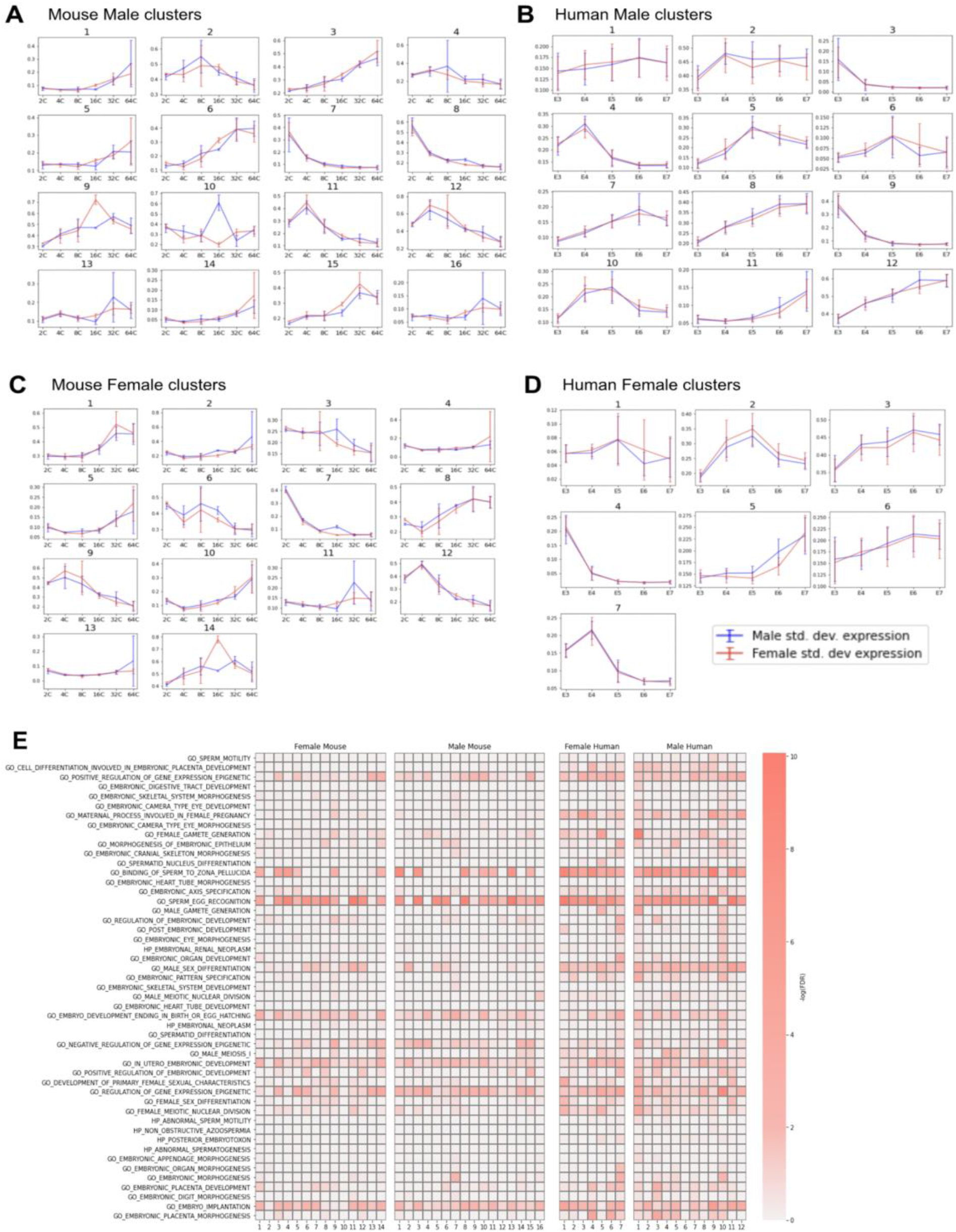
Derived sex-biased genes and networks between mouse and human. A - D. Clusters of similar expression resulting from non-negative matrix factorization of expression data. Each plot depicts mean expression across samples for each stage in (A) male mouse clusters, (B) male human clusters, (C) female mouse clusters, and (D) female human clusters. (E) Heatmap of enriched pathways across NMF clusters.

A heatmap of GSEA of NMF clusters shows concordance between enriched clusters in mouse and human that include signals from male and female gametes, possibly remnants of events during fertilization (Figure 6E) (Supplementary Figures S13-17).

## DISCUSSION

In this study, we observe molecular signals of sexual identity during the earliest stages in mammalian embryogenesis, during the first cell divisions immediately after fertilization. These observations challenge the prevailing view that sexual differentiation occurs only after SRY activates gonad formation and subsequent circulation of sex-specific hormones. From single-cell transcriptomics results in both mice and humans, we observe the establishment of rapidly changing male and female network landscapes that are enriched in X-linked protein coding genes involved in sex-specific cellular development (Table 1). Significant differences exist in gene ontological processes among male and female cells during early stages of development. In female cells, biological processes belonging to cell organization and cell compartment are delayed, while in male cells, RNA and DNA related processes are delayed. Overall, our results reveal the emergence of early-acting sexual networks among males and females, thus, placing prominence on the role of sex chromosomes during early stages of organismal development^34,35^.

Our work also highlights early developmental differences across mammals. Early stages of embryogenesis are generally characterized by a high degree of morphological conservation across mammals which has been historically associated to recapitulation^36,37^. There are several known differences in preimplantation embryogenesis between mice and humans. Human early development is protracted compared to mouse, with zygotic genome activation at the 8-cell stage, whereas in mouse it occurs at the 2-cell stage. Maternal transcripts exhibit differences also, with human maternal programs extended until the later zygotic genome activation.

However, recent decades of molecular studies have shown that the underlying expression programs between overtly similar processes differ significantly between humans and mice in both content and timing^38,39^. For example, some of the master regulators that determine the fates of the outer trophectoderm and the inner cell mass act through different pathways in mice and humans or are different altogether^21,38^. With new technologies, we can now identify critical milestones such as the zygotic genome activation (ZGA)^40,41^ and the first and second lineage segregations that give rise to the future trophectoderm, epiblast, and primitive endoderm. The availability of single-cell sequencing data allows us to interrogate sex-specific gene expression across the earliest stages of development.

X-chromosome inactivation in mouse and human female cells also exhibit different dynamics. In mice, the paternal X chromosome is preferentially inactivated, and then reactivated during pre-implantation, with random X chromosome inactivation occurring upon implantation^42^. In humans both maternal and paternal X chromosomes are initially active, despite biallelic expression of *Xist*, with random inactivation initiating and progressing throughout pre-implantation^17^.

Several regulatory factors exhibit sex-biased expression with different dynamics in male and female mouse embryos. For example, *Rfx4, Hnf1b, Glis3, Sox17* and *Smad9* are male-biased at earlier stages and female-biased later. On the other hand, *Hey1, Hoxb13, Setd7, Irf5* and *Hif1a* are female-biased at earlier stages and male-biased later. This suggests that, at least for processes regulated by these factors, pre-implantation development is offset between males and females. However, a substantial number of regulatory factors show sex-specific expression throughout this phase, the majority of which are encoded by autosomal genes. *Kdm5d*, a Y-linked histone lysine demethylase, is expressed throughout all stages in the males, whereas its X-linked homologue, *Kdm6a* is female-biased only at the 8- and 16-cell stages, either because the paternal X is undergoing reactivation or because it escapes paternal X inactivation. Interestingly, both the mouse and human embryos exhibit a diminishing of sex-biased expression as development progresses towards formation of the blastocyst and that the male embryos exhibit more Y-linked genes than the females do for X-linked genes. We hypothesize that some of the initial sex differences in expression could reflect that the reprogramming of the Y chromosome, namely, the replacement of the protamines, could occur faster than that of the X chromosome provided by the sperm, due to their size differences. This would agree with the greater Y-linked expression in the male embryos.

Our study’s conclusions have direct implications on the study of sexual differences across later stages of development and on sex-specific disease^43^. The sex-biased expression of transcriptional and epigenetic factors reported here prompts questions on how sex-specific networks in pre-implantation affect sex biased gene expression after implantation, as lineage determination and organogenesis proceed, including how hormones interact with pre-established sex differences by compensating for or enhancing them. We also note that since transcription factors are generally expressed at low levels, many sex-biased genes may be underrepresented in single-cell RNA-seq experiments. In addition, relative to our much larger set of human samples, the sample sizes for the earliest stages of mouse embryogenesis are small (Supplementary Figure 1) and additional studies would increase the power to detect sex-biased expression. Furthermore, earlier time points for human embryos would allow a more precise match of the developmental stages with the mouse. Thus, we anticipate that there are more differences in gene expression of regulatory factors than detected in our study. In addition, with the increasing availability of single-cell RNA-seq data from specific cell types in embryos, future studies will allow more detailed comparisons between mouse and human to find common substructures, facilitating the discovery of conserved functional modules. Future studies on the development of sex-specific networks activated by sex chromosome differences in very early embryogenesis will be able address these questions with greater precision.

## ACKNOWLEDGEMENTS

This manuscript was funded by NSF #1933738 to NE and RJK.

## SUPPLEMENTARY INFORMATION

### Supplementary Tables

**Supplementary Table S1**.

Transcriptomic datasets used across developmental time and sex. (page 2).

**Supplementary Table S2**.

Differentially expressed genes in mice (page 3).

**Supplementary Table S3**.

Differentially expressed genes in humans (page 52).

**Supplementary Table S4**.

Male mice: Gene set enrichment analysis results of enriched pathways across NMF clusters (page 148).

**Supplementary Table S5**.

Female mice: Gene set enrichment analysis results of enriched pathways across NMF clusters (page 283).

**Supplementary Table S6**.

Male humans: Gene set enrichment analysis results of enriched pathways across NMF clusters (page 343).

**Supplementary Table S7**.

Female humans: Gene set enrichment analysis results of enriched pathways across NMF clusters (page 426).

**Supplementary Table S8**.

Male mice: NMF coefficient matrix of gene expression data (page 557).

**Supplementary Table S9**.

Female mice: NMF coefficient matrix of gene expression data (page 787).

**Supplementary Table S10**.

Male humans: NMF coefficient matrix of gene expression data (page 1046).

**Supplementary Table S11**.

Female humans: NMF coefficient matrix of gene expression data (page 1500).

### Supplementary Figures

**Supplementary Figure S1**.

Distributions of counts per gene for unfiltered and filtered expression data in mice and humans.

**Supplementary Figure S2**.

Genes with the highest and lowest principal component scores for the top 30 principal components of gene expression data in mice.

**Supplementary Figure S3**.

Genes with the highest and lowest principal component scores for the top 30 principal components of gene expression data in humans.

**Supplementary Figure S4**.

Median number of enriched NMF clusters per biological process GOSlim term, normalized by cluster size, between mouse male and female.

**Supplementary Figure S5**.

Median number of enriched NMF clusters per biological process GOSlim term, normalized by cluster size, between human male and female.

**Supplementary Figure S6**.

Median number of enriched NMF clusters per cellular component GOSlim term, normalized by cluster size, between mouse male and female.

**Supplementary Figure S7**.

Median number of enriched NMF clusters per cellular component GOSlim term, normalized by cluster size, between human male and female.

**Supplementary Figure S8**.

Median number of enriched NMF clusters per molecular function GOSlim term, normalized by cluster size, between mouse male and female.

**Supplementary Figure S9**.

Median number of enriched NMF clusters per cellular component GOSlim term, normalized by cluster size, between human male and female.

**Supplementary Figure S10**.

Scatter plot of the median number of enriched NMF clusters per biological process GOSlim term, normalized by cluster size, between (top) Male mouse vs. human and (bottom) Female mouse vs. human. Top 5 terms with the largest difference between mouse and human are labeled.

**Supplementary Figure S11**.

Scatter plot of the median number of enriched NMF clusters per cellular component GOSlim term, normalized by cluster size, between (top) Male mouse vs. human and (bottom) Female mouse vs. human. Top 5 terms with the largest difference between mouse and human are labeled.

**Supplementary Figure S12**.

Scatter plot of the median number of enriched NMF clusters per molecular function GOSlim term, normalized by cluster size, between (top) Male mouse vs. human and (bottom) Female mouse vs. human. Top 5 terms with the largest difference between mouse and human are labeled.

**Supplementary Figure S13**.

Heatmap of the normalized number of NMF clusters enriched under each GOslim term for biological process.

**Supplementary Figure S14**.

Heatmap of the normalized number of NMF clusters enriched under each GOslim term for cellular component.

**Supplementary Figure S15**.

Heatmap of the normalized number of NMF clusters enriched under each GOslim term for molecular function.

**Supplementary Figure S16**.

Average expression value of each gene across samples in male and female mice. Each plot indicates a cluster (metagene), and plot titles indicate cluster size. Green lines indicate embryonic stage.

**Supplementary Figure S16**.

Average expression value of each gene across samples in male and female humans. Each plot indicates a cluster (metagene), and plot titles indicate cluster size. Green lines indicate embryonic stage.

