## Supplementary Figures for "Comparative developmental genomics of sex-biased gene expression in early embryogenesis across mammals"

**Supplementary Figure S1.** Distributions of counts per gene for unfiltered and filtered expression data in mice and humans.

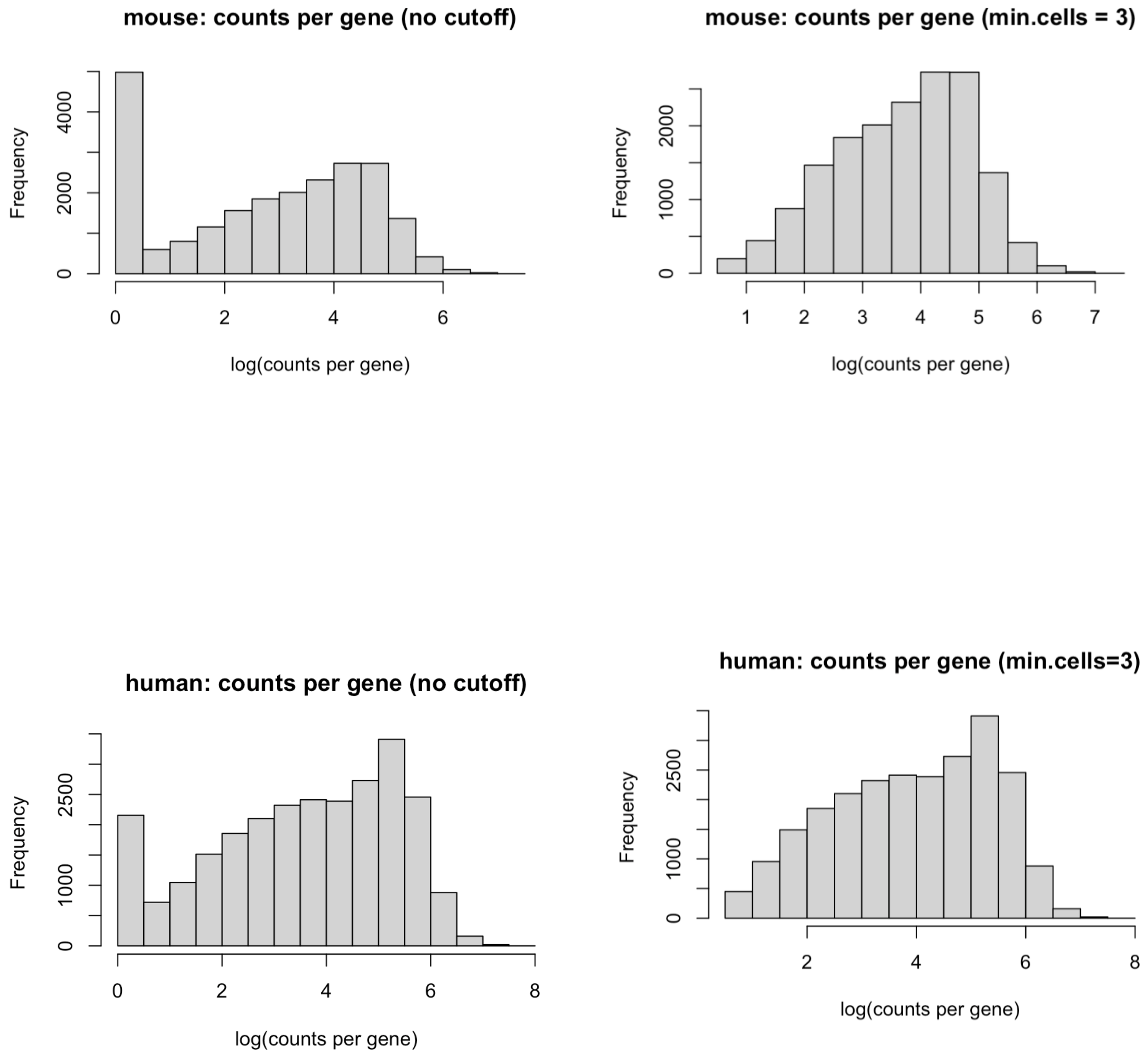

**Supplementary Figure S2.** Genes with the highest and lowest principal component scores for the top 30 principal components of gene expression data in mice.

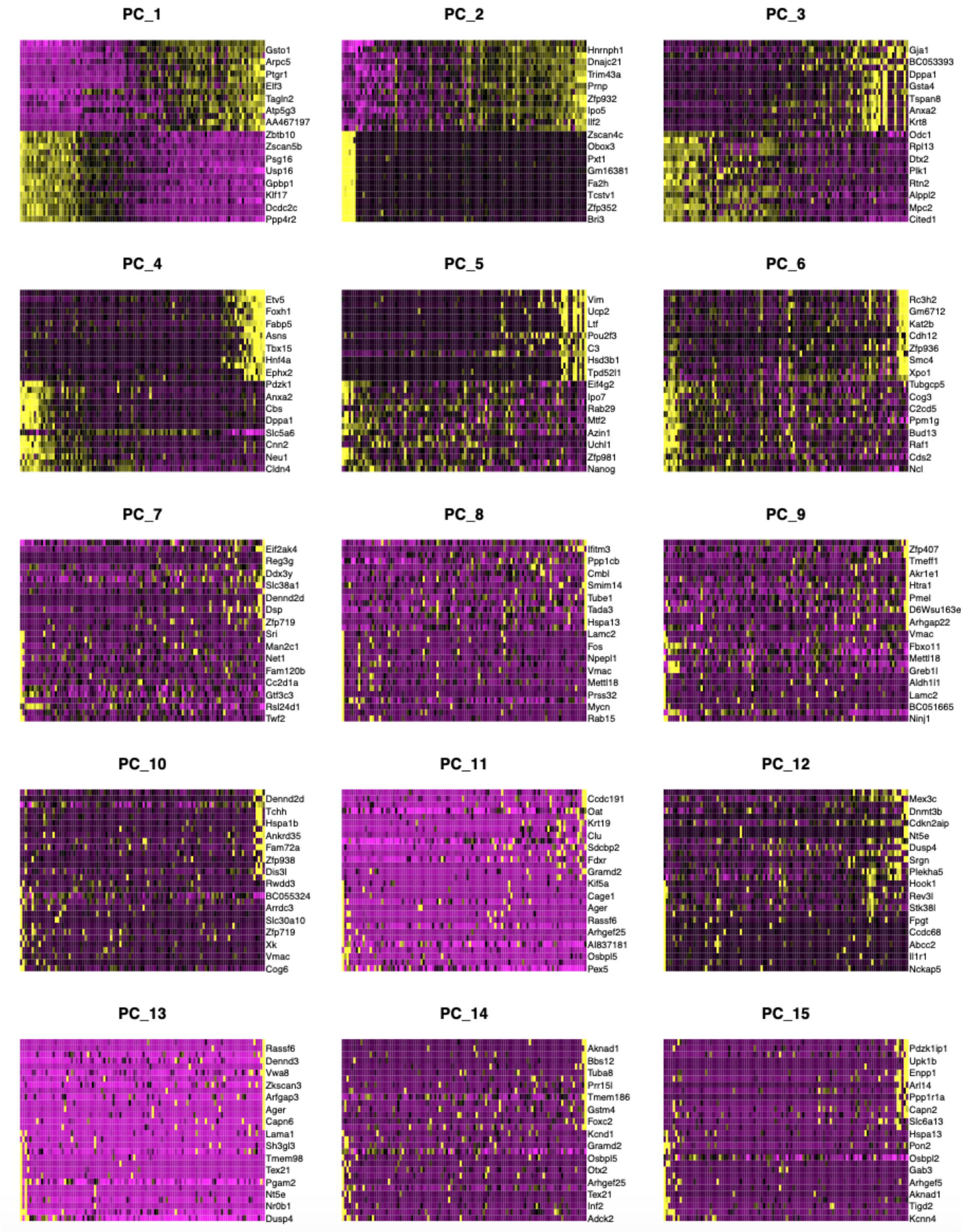

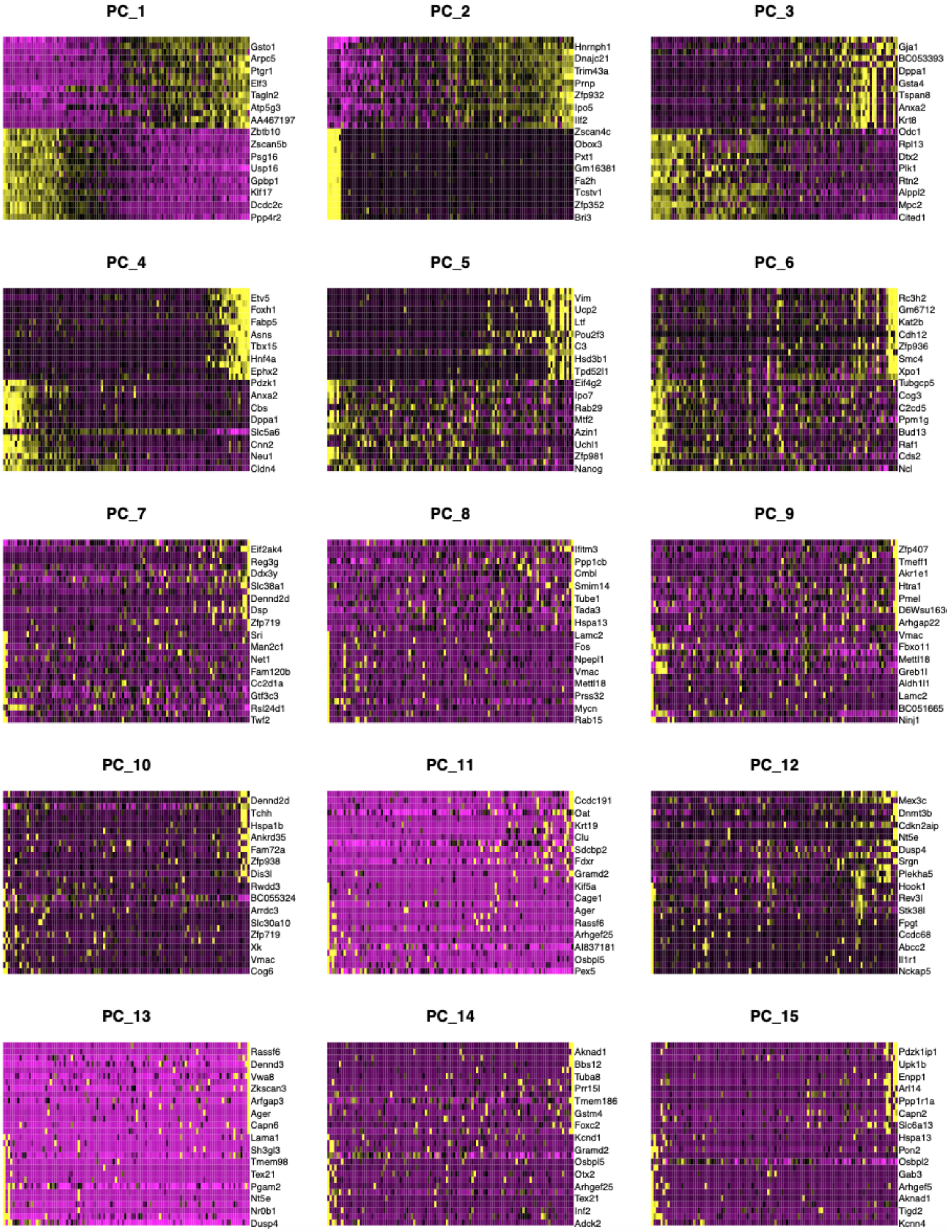
PC_1
Gsto1
Arpc5
Ptgr1
Elf3
Tagln2
Atp5g3
AA467197
Zbtb10
Zscan5b
Psg16
Usp16
Gpbp1
Klf17
Dcdc2c
Ppp4r2
PC_2
Hnrnph1
Dnajc21
Trim43a
Prnp
Zfp932
Ipo5
Ilf2
Zscan4c
Obox3
Pxt1
Gm16381
Fa2h
Tcstv1
Zfp352
Bri3
PC_3
Gja1
BC053393
Dppa1
Gsta4
Tspan8
Anxa2
Krt8
Odc1
Rpl13
Dtx2
Plk1
Rtn2
Alppl2
Mpc2
Cited1
PC_4
Etv5
Foxh1
Fabp5
Asns
Tbx15
Hnf4a
Ephx2
Pdzk1
Anxa2
Cbs
Dppa1
Slc5a6
Cnn2
Neu1
Cldn4
PC_5
Vim
Ucp2
Ltf
Pou2f3
C3
Hsd3b1
Tpd52l1
Eif4g2
Ipo7
Rab29
Mtf2
Azin1
Uchl1
Zfp981
Nanog
PC_6
Rc3h2
Gm6712
Kat2b
Cdh12
Zfp936
Smc4
Xpo1
Tubgcp5
Cog3
C2cd5
Ppm1g
Bud13
Raf1
Cds2
Ncl
PC_7
Eif2ak4
Reg3g
Ddx3y
Slc38a1
Dennd2d
Dsp
Zfp719
Sri
Man2c1
Net1
Fam120b
Cc2d1a
Gtf3c3
Rsl24d1
Twf2
PC_8
Ifitm3
Ppp1cb
Cmbl
Smim14
Tube1
Tada3
Hspa13
Lamc2
Fos
Npepl1
Vmac
Mettl18
Prss32
Mycn
Rab15
PC_9
Zfp407
Tmeff1
Akr1e1
Htra1
Pmel
D6Wsu163
Arhgap22
Vmac
Fbxo11
Mettl18
Greb1l
Aldh1l1
Lamc2
BC051665
Ninj1
PC_10
Dennd2d
Tchh
Hspa1b
Ankrd35
Fam72a
Zfp938
Dis3l
Rwdd3
BC055324
Arrdc3
Slc30a10
Zfp719
Xk
Vmac
Cog6
PC_11
Ccdc191
Oat
Krt19
Clu
Sdcbp2
Fdxr
Gramd2
Kif5a
Cage1
Ager
Rassf6
Arhgef25
AI837181
Osbpl5
Pex5
PC_12
Mex3c
Dnmt3b
Cdkn2aip
Nt5e
Dusp4
Srgn
Plekha5
Hook1
Rev3l
Stk38l
Fpgt
Ccdc68
Abcc2
Il1r1
Nckap5
PC_13
Rassf6
Dennd3
Vwa8
Zkscan3
Arfgap3
Ager
Capn6
Lama1
Sh3gl3
Tmem98
Tex21
Pgam2
Nt5e
Nr0b1
Dusp4
PC_14
Aknad1
Bbs12
Tuba8
Prr15l
Tmem186
Gstm4
Foxc2
Kcnd1
Gramd2
Osbpl5
Otx2
Arhgef25
Tex21
Inf2
Adck2
PC_15
Pdzk1ip1
Upk1b
Enpp1
Arl14
Ppp1r1a
Capn2
Slc6a13
Hspa13
Pon2
Osbpl2
Gab3
Arhgef5
Aknad1
Tigd2
Kcnn4

**Supplementary Figure S3.** Genes with the highest and lowest principal component scores for the top 30 principal components of gene expression data in humans.

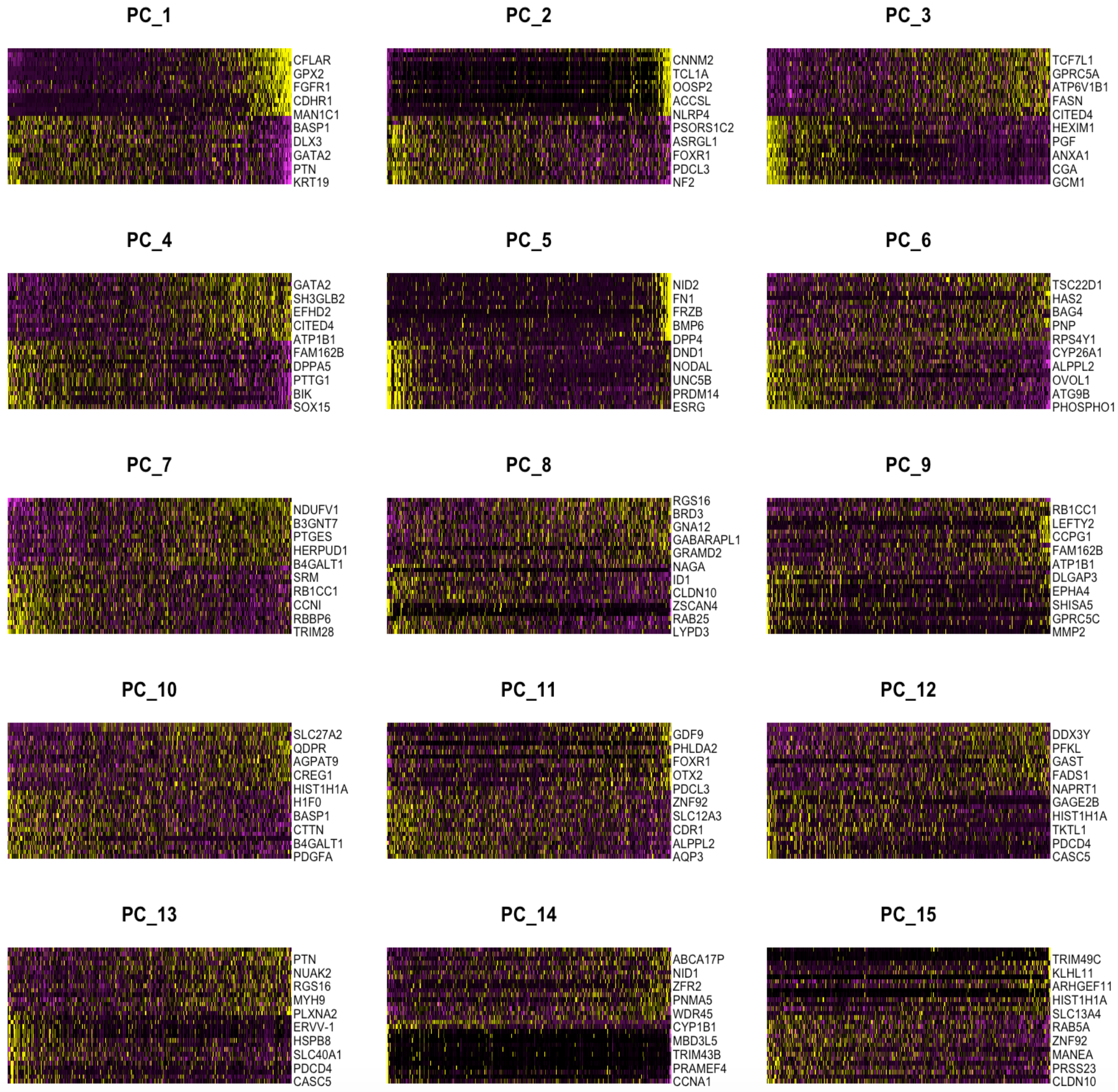

**Supplementary Figure S4.** Median number of enriched NMF clusters per biological process GOSlim term, normalized by cluster size, between mouse male and female.

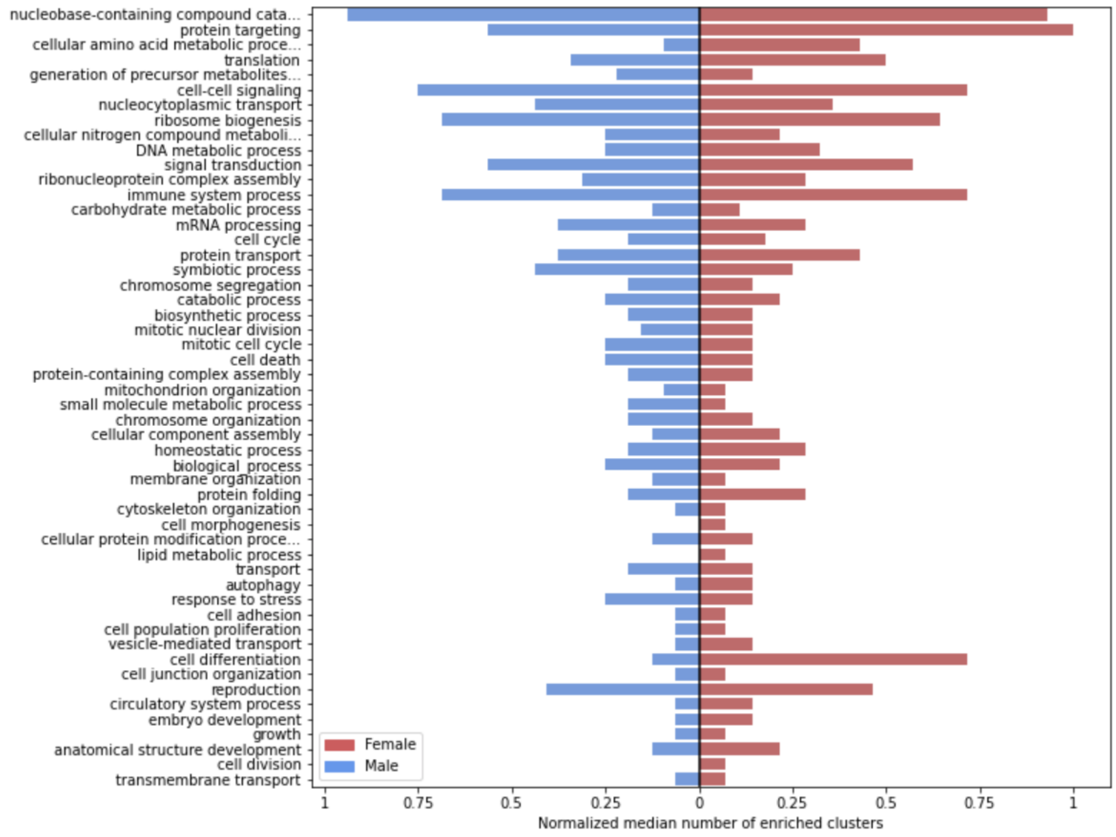

**Supplementary Figure S5.** Median number of enriched NMF clusters per biological process GOSlim term, normalized by cluster size, between human male and female

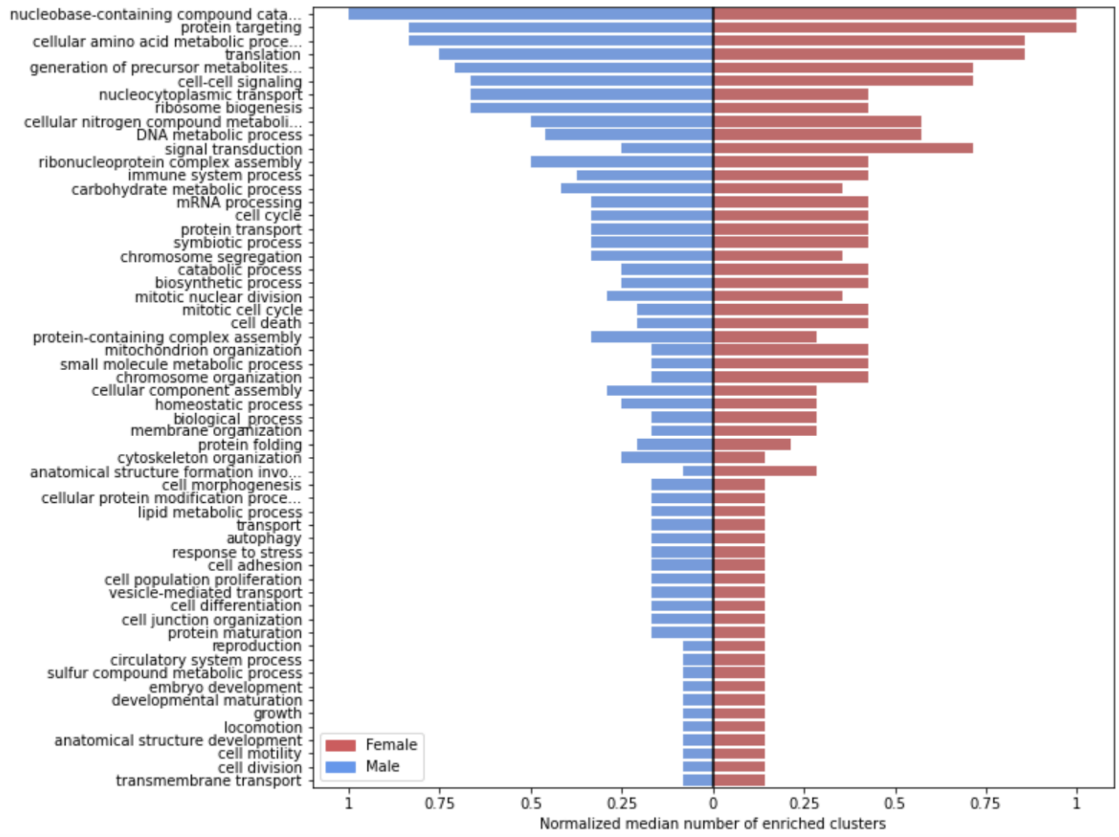


.

**Supplementary Figure S6.** Median number of enriched NMF clusters per cellular component GOSlim term, normalized by cluster size, between mouse male and female

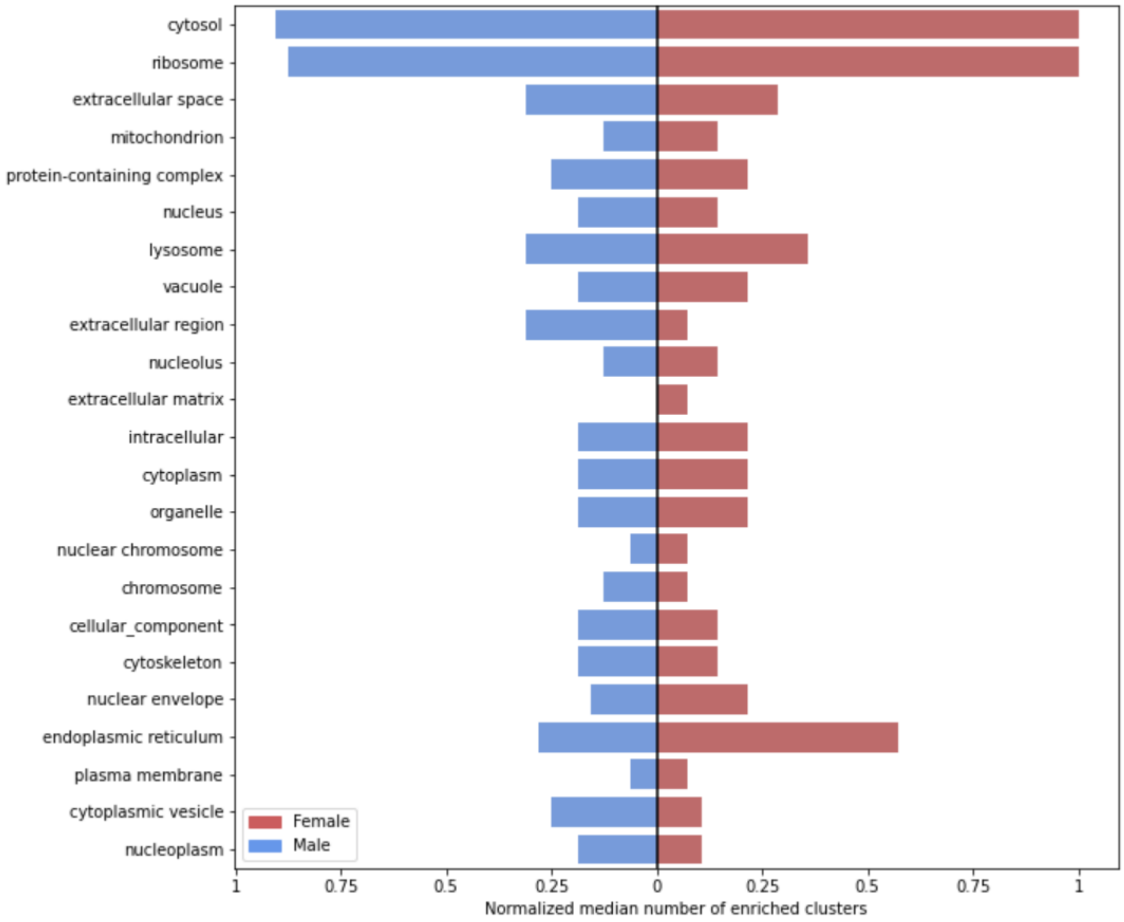


.

**Supplementary Figure S7.** Median number of enriched NMF clusters per cellular component GOSlim term, normalized by cluster size, between human male and female.

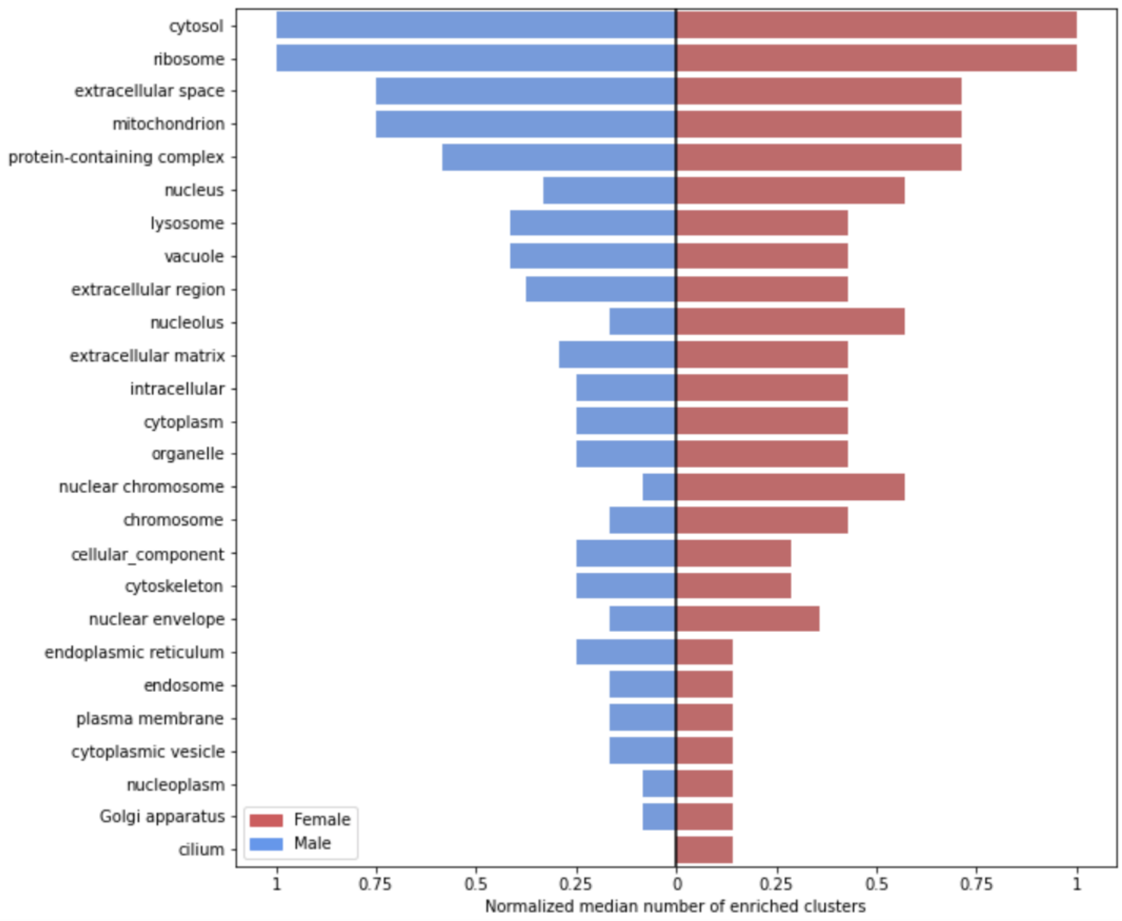

**Supplementary Figure S8.** Median number of enriched NMF clusters per molecular function. GOSlim term, normalized by cluster size, between mouse male and female

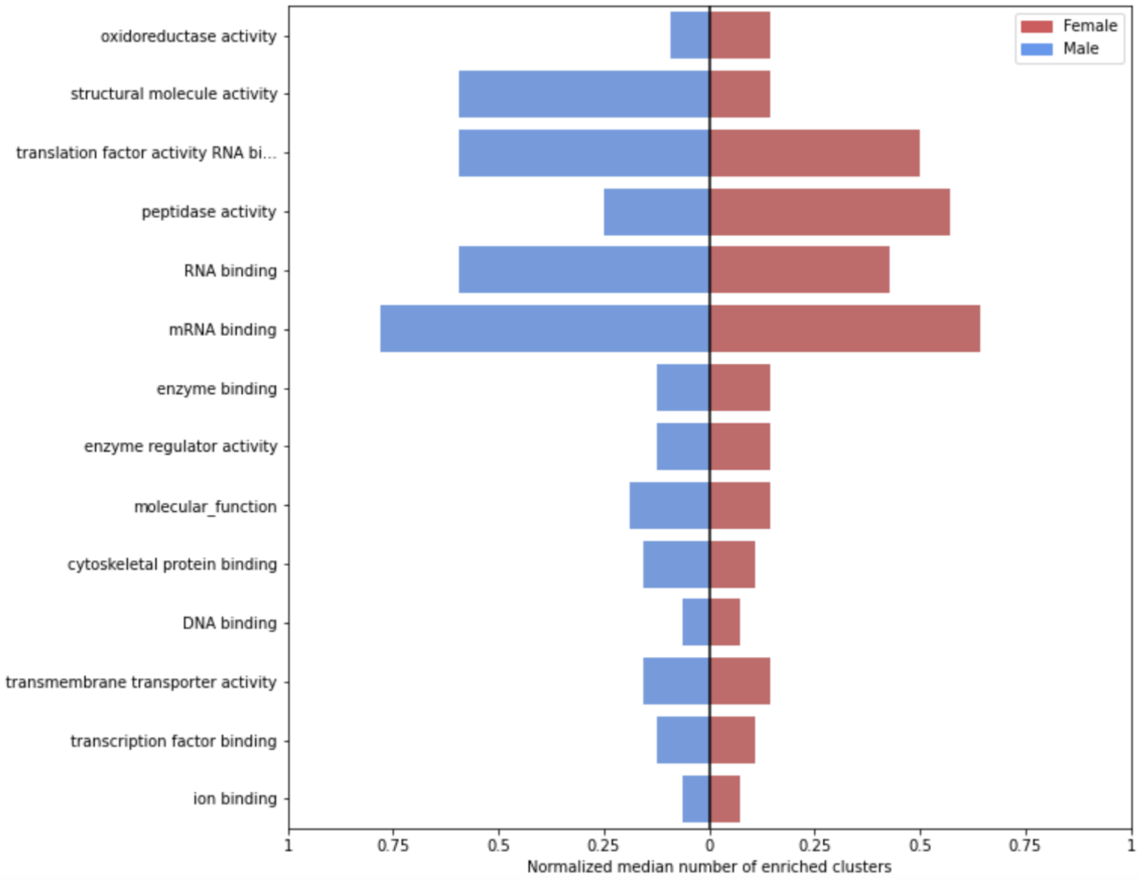

**Supplementary Figure S9.** Median number of enriched NMF clusters per cellular component GOSlim term, normalized by cluster size, between human male and female.

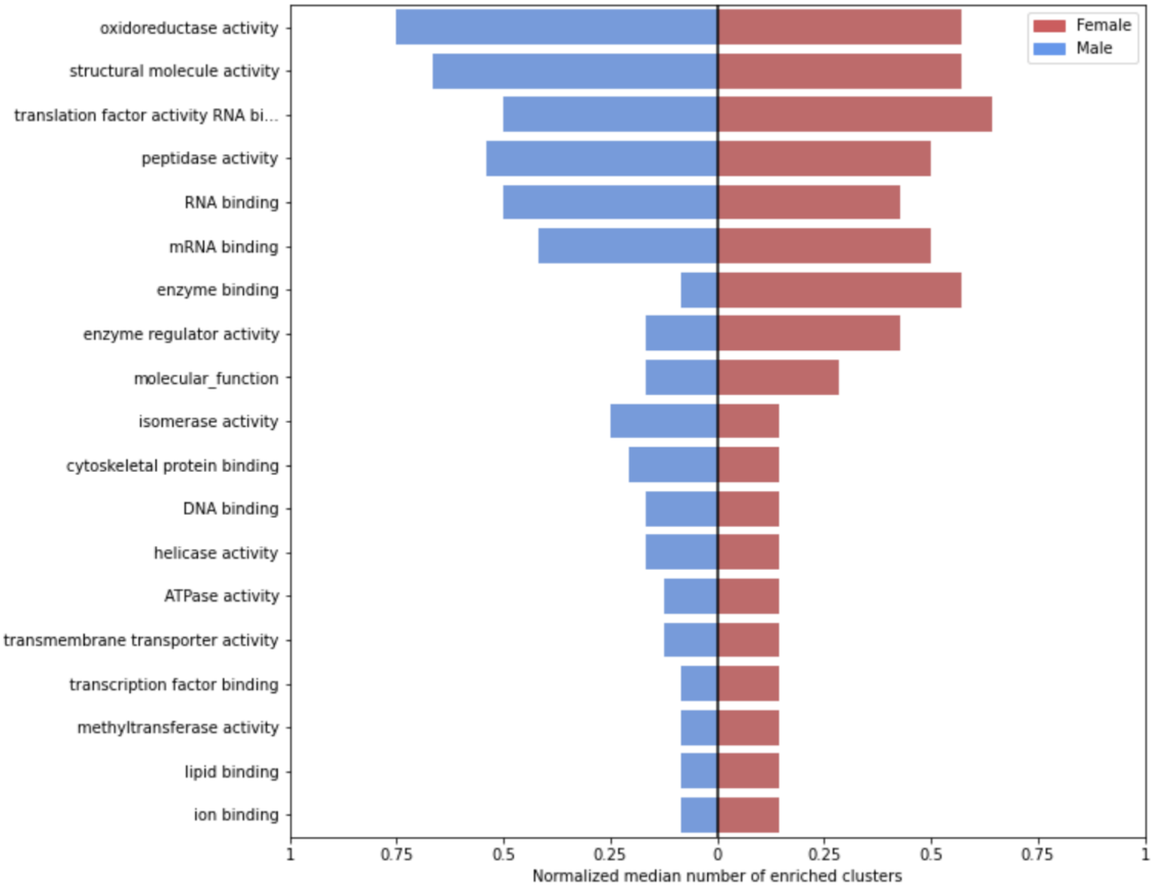

**Supplementary Figure S10.** Median number of enriched NMF clusters per biological process GOSlim term, normalized by cluster size, between (top) Male mouse vs. human and (bottom) Female mouse vs. human. Top 5 terms with the largest difference between mouse and human are labeled.

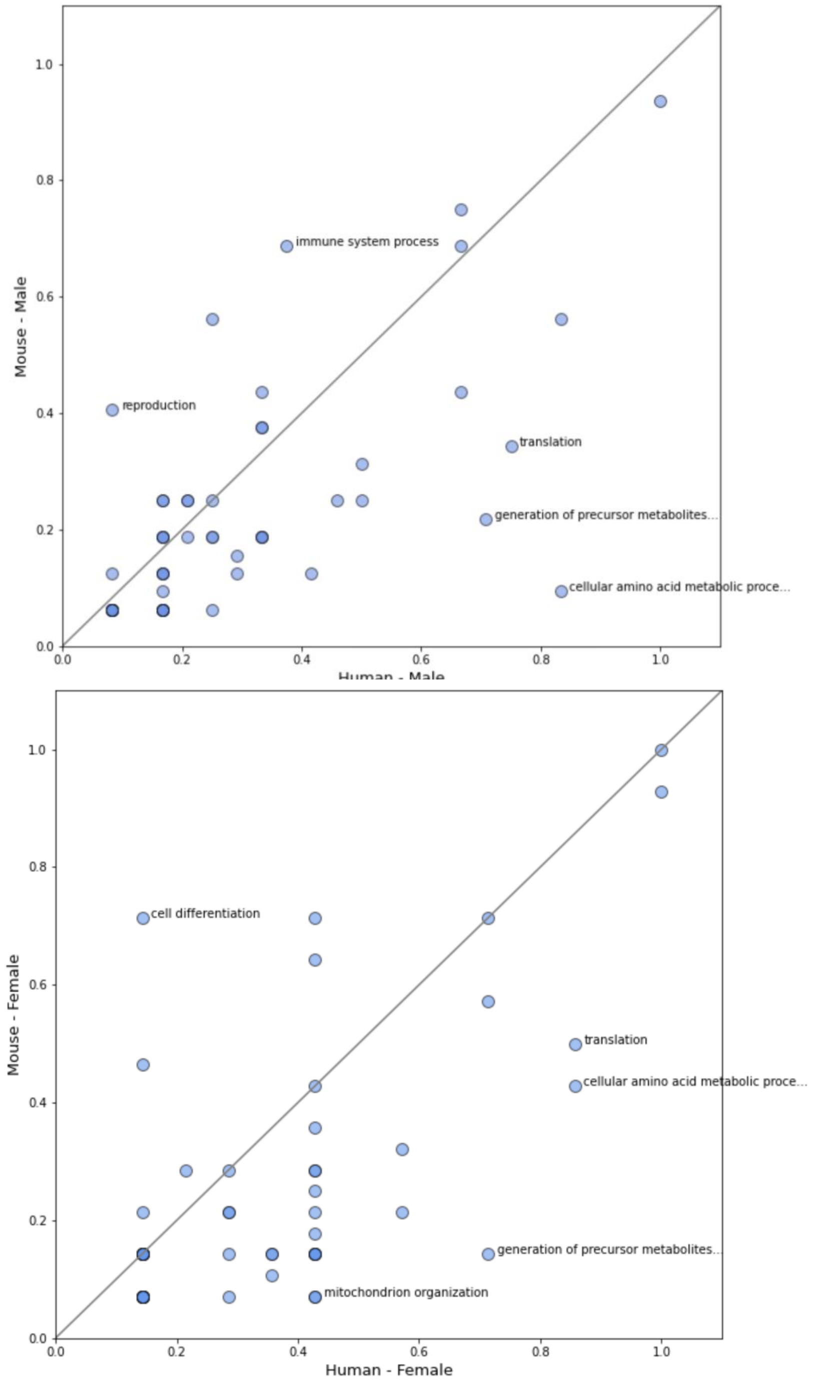

**Supplementary Figure S11.** Median number of enriched NMF clusters per cellular component GOSlim term, normalized by cluster size, between (top) Male mouse vs. human and (bottom) Female mouse vs. human. Top 5 terms with the largest difference between mouse and human are labeled

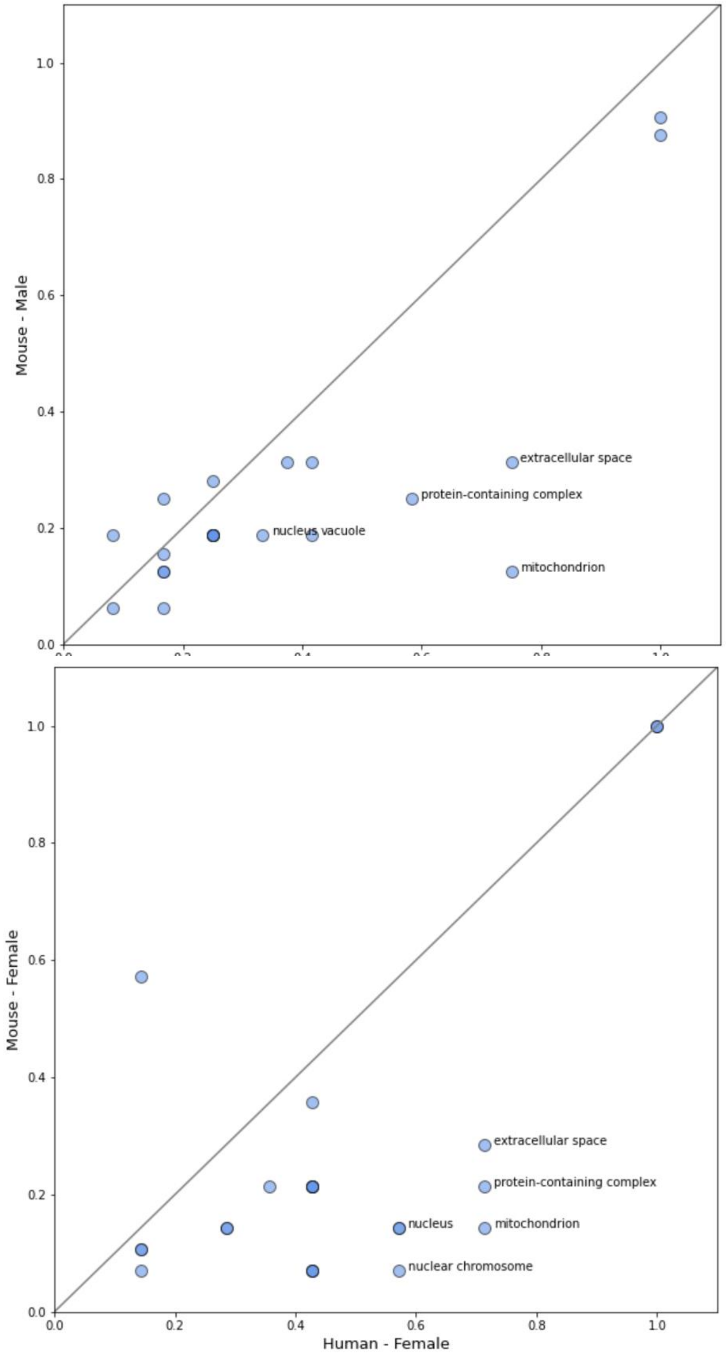


.

**Supplementary Figure S12.** Median number of enriched NMF clusters per molecular function GOSlim term, normalized by cluster size, between (top) Male mouse vs. human and (bottom) Female mouse vs. human. Top 5 terms with the largest difference between mouse and human are labeled.

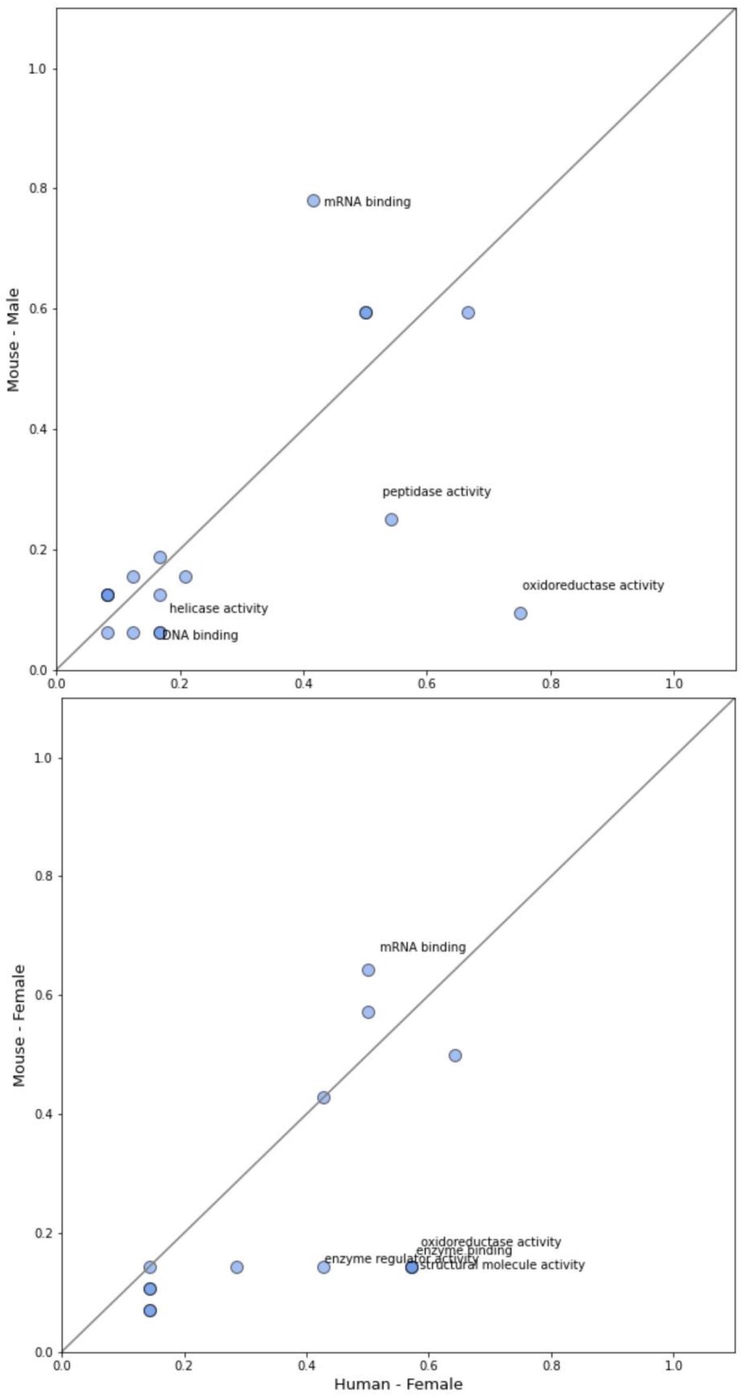

**Supplementary Figure S13.** Heatmap of the normalized number of NMF clusters enriched under each GOslim term for biological process.

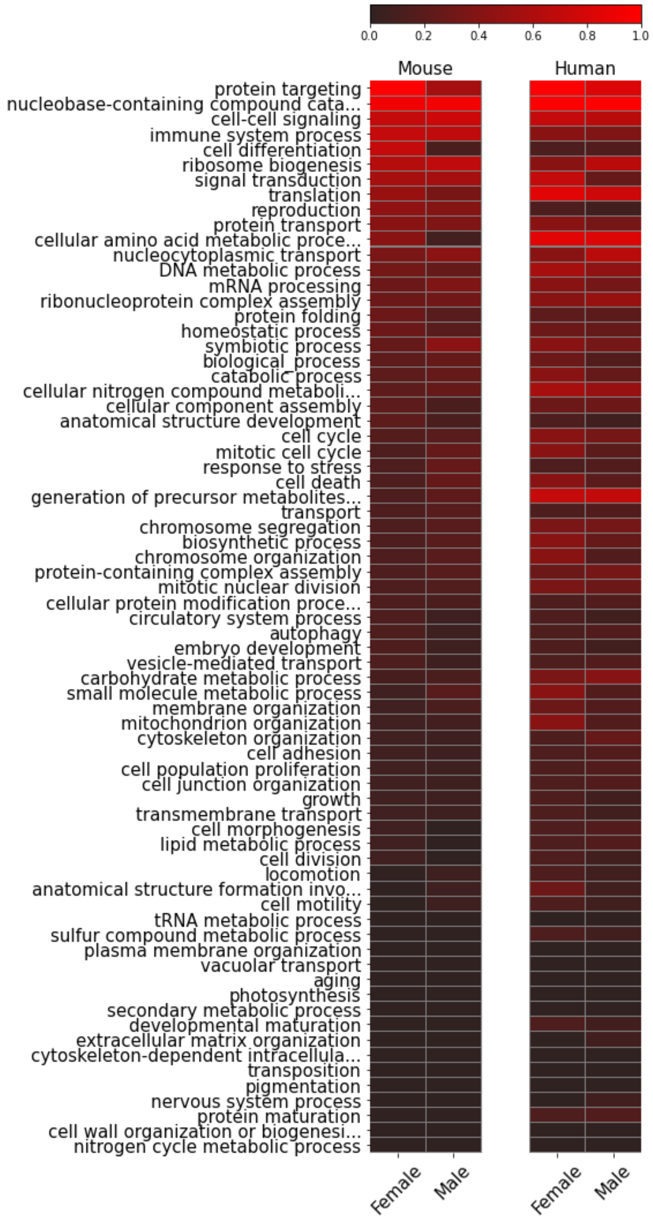

**Supplementary Figure S14.** Heatmap of the normalized number of NMF clusters enriched under each GOslim term for cellular component.

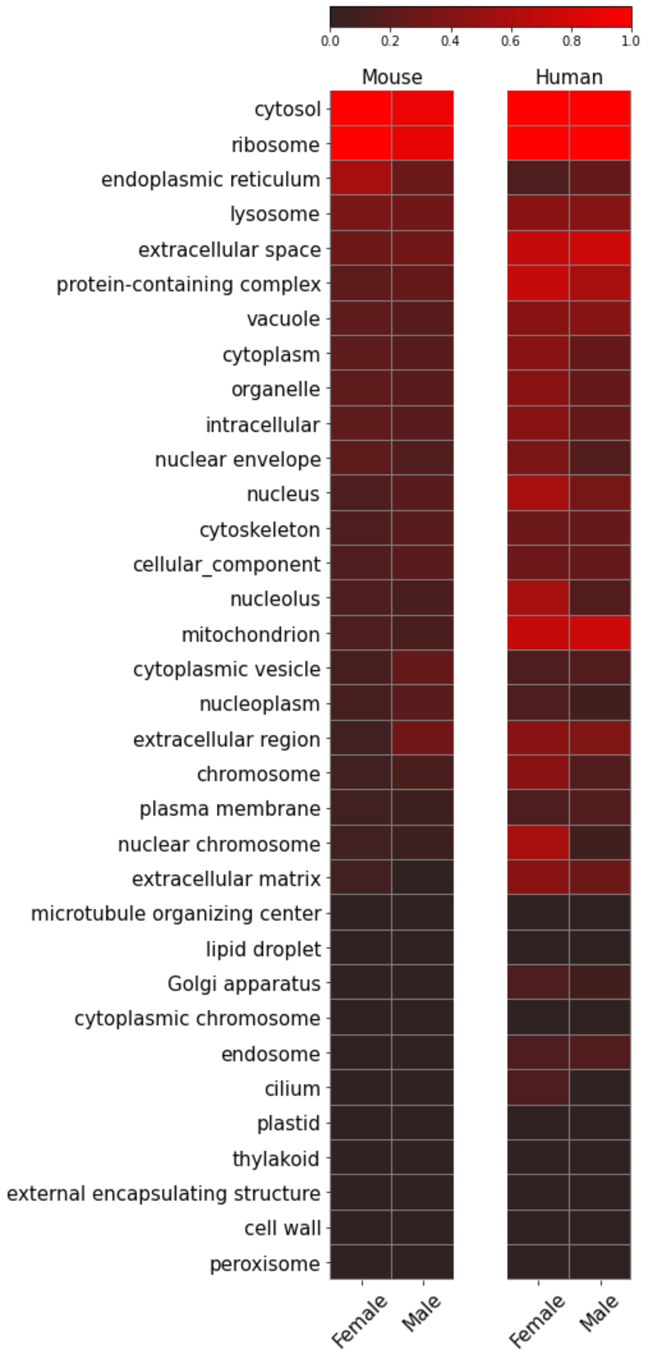

**Supplementary Figure S15.** Heatmap of the normalized number of NMF clusters enriched under each GOslim term for molecular function.

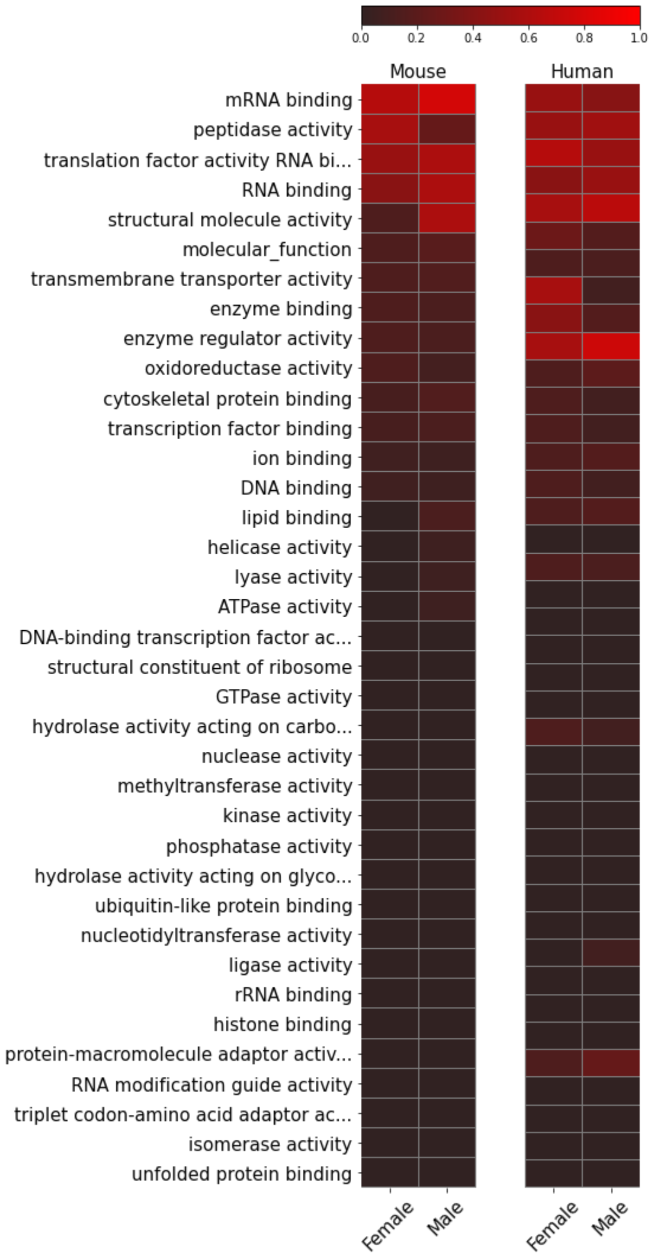

**Supplementary Figure S16.** Average expression value of each gene across samples in male and female mice. Each plot indicates a cluster (metagene), and plot titles indicate cluster size. Green lines indicate embryonic stage.

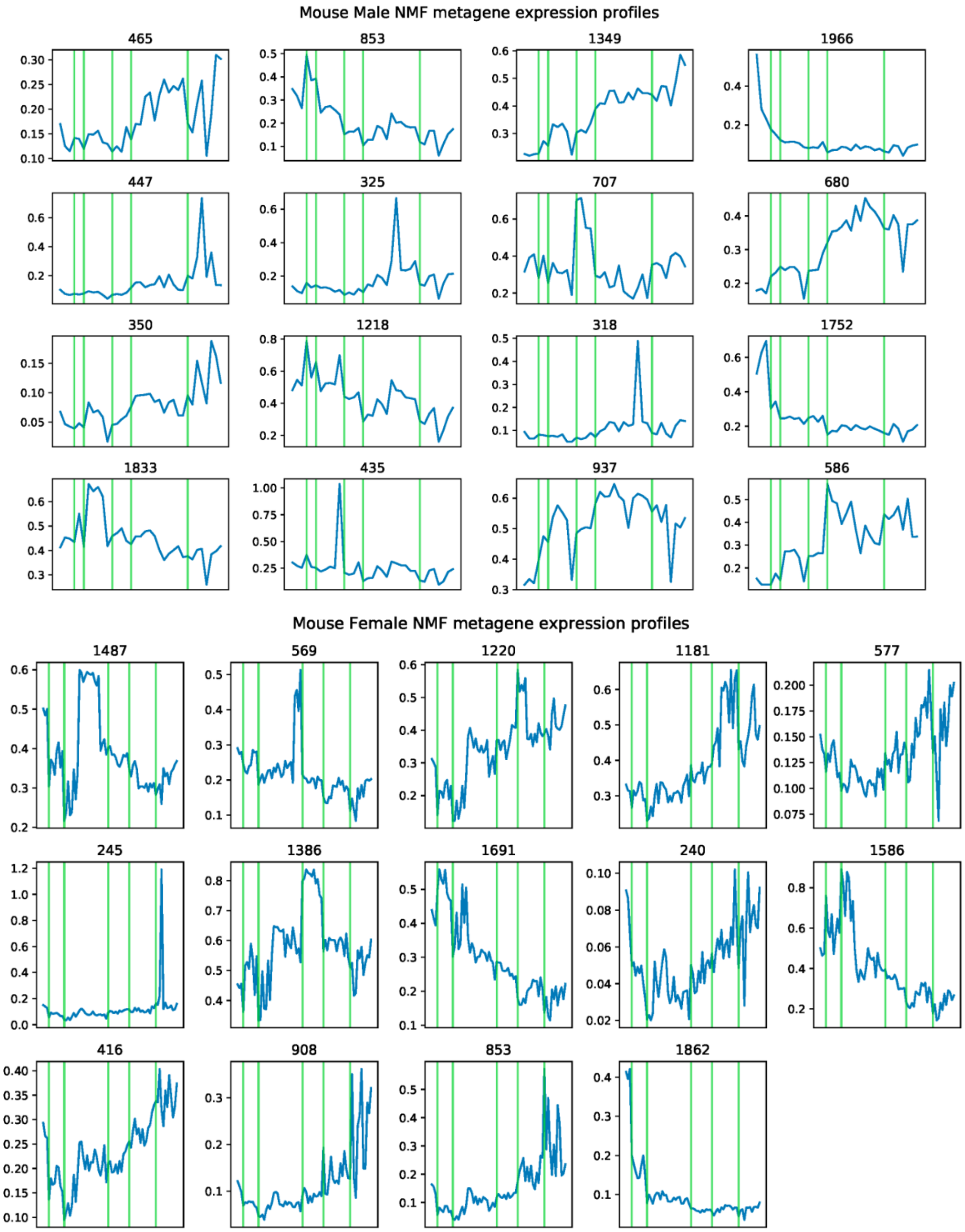

**Supplementary Figure S17.** Average expression value of each gene across samples in male and female humans. Each plot indicates a cluster (metagene), and plot titles indicate cluster size. Green lines indicate embryonic stage.

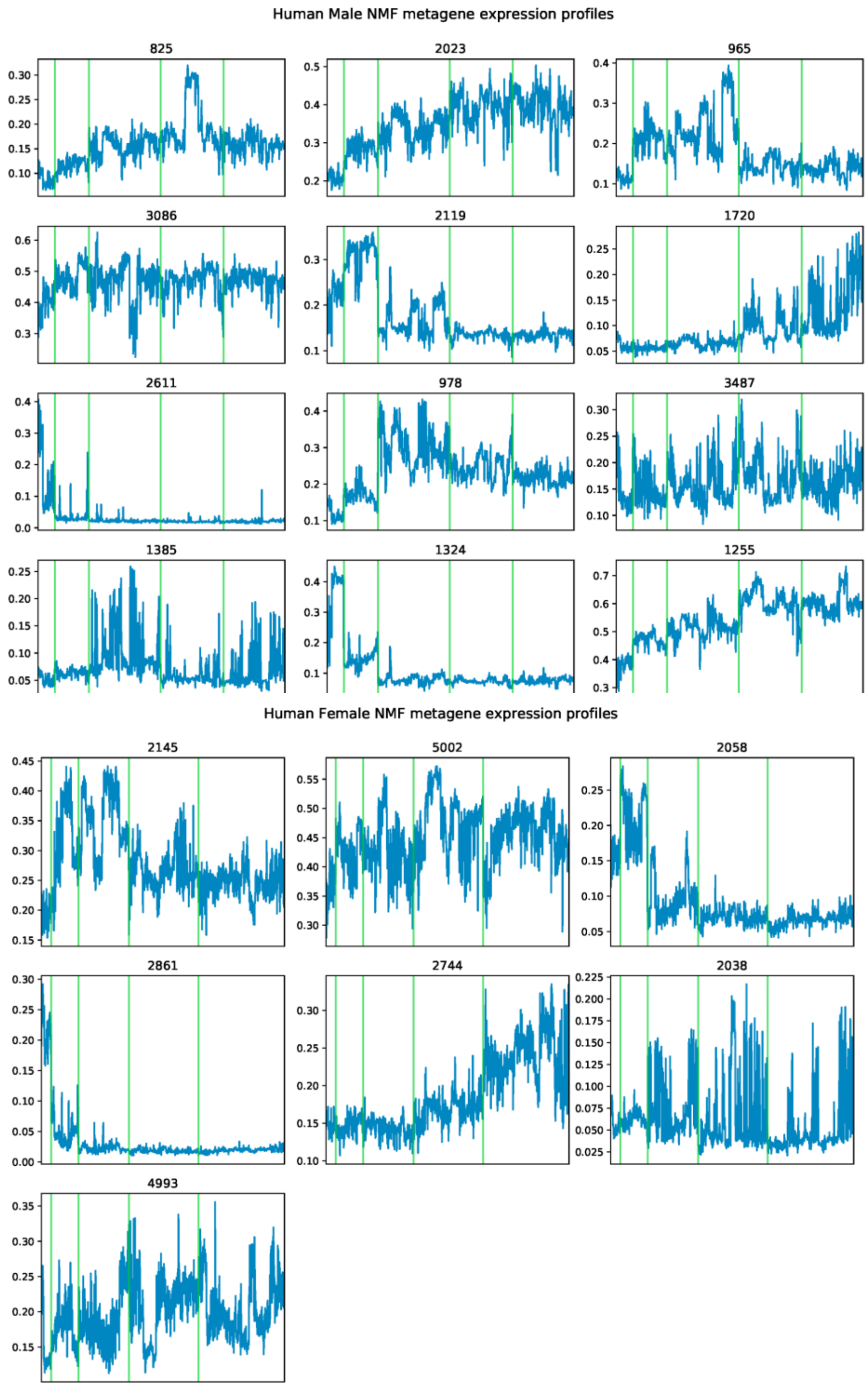
